# Systemic inflammation and lymphocyte activation precede rheumatoid arthritis

**DOI:** 10.1101/2024.10.25.620344

**Authors:** Ziyuan He, Marla C. Glass, Pravina Venkatesan, Marie L. Feser, Leander Lazaro, Lauren Y. Okada, Nhung T. T. Tran, Yudong D. He, Samir Rachid Zaim, Christy E. Bennett, Padmapriyadarshini Ravisankar, Elisabeth M. Dornisch, Najeeb A. Arishi, Ashley G. Asamoah, Saman Barzideh, Lynne A. Becker, Elizabeth A. Bemis, Jane H. Buckner, Christopher E. Collora, Megan A. L. Criley, M. Kristen Demoruelle, Chelsie L. Fleischer, Jessica Garber, Palak C. Genge, Qiuyu Gong, Lucas T. Graybuck, Claire E. Gustafson, Brian C. Hattel, Veronica Hernandez, Alexander T. Heubeck, Erin K. Kawelo, Upaasana Krishnan, Emma L. Kuan, Kristine A. Kuhn, Christian M. LaFrance, Kevin J. Lee, Ruoxin Li, Cara Lord, Regina R. Mettey, Laura Moss, Blessing Musgrove, Kathryn Nguyen, Andrea Ochoa, Vaishnavi Parthasarathy, Mark-Phillip Pebworth, Chong Pedrick, Tao Peng, Cole G. Phalen, Julian Reading, Charles R. Roll, Jennifer A. Seifert, Marguerite D. Siedschlag, Cate Speake, Christopher C. Striebich, Tyanna J. Stuckey, Elliott G. Swanson, Hideto Takada, Tylor Thai, Zachary J. Thomson, Nguyen Trieu, Vlad Tsaltskan, Wei Wang, Morgan D. A. Weiss, Amy Westermann, Fan Zhang, David L. Boyle, Ananda W. Goldrath, Thomas F. Bumol, Xiao-jun Li, V. Michael Holers, Peter J. Skene, Adam K. Savage, Gary S. Firestein, Kevin D. Deane, Troy R. Torgerson, Mark A. Gillespie

**Author notes:** Corresponding authors. (KDD); (TRT); (MAG). These authors contributed equally to this work (Ziyuan He, Marla C. Glass). These authors contributed equally to this work (Adam K. Savage, Gary S. Firestein, Kevin D. Deane, Troy R. Torgerson, Mark A. Gillespie).

## Abstract

Some autoimmune diseases, including rheumatoid arthritis (RA), are preceded by a critical subclinical phase of disease activity. Proactive clinical management is hampered by a lack of biological understanding of this subclinical ‘at-risk’ state and the changes underlying disease development. In a cross-sectional and longitudinal multi-omics study of peripheral immunity in the autoantibody-positive at-risk for RA period, we identified systemic inflammation, proinflammatory-skewed B cells, expanded Tfh17-like cells, epigenetic bias in naive T cells, TNF+IL1B+ monocytes resembling a synovial macrophage population, and CD4 T cell transcriptional features resembling those suppressed by abatacept (CTLA4-Ig) in RA patients. Our findings characterize pathogenesis prior to clinical diagnosis and suggest the at-risk state exhibits substantial immune alterations that could potentially be targeted for early intervention to delay or prevent autoimmunity. We provide a suite of tools at https://apps.allenimmunology.org/aifi/insights/ra-progression/ to facilitate exploration and enhance accessibility of this extensive dataset.

**One Sentence Summary:** ACPA+ at-risk individuals show RA-like inflammation and multi-compartment immune dysregulation during transition to clinically active RA

## Main Text

In autoimmunity, early diagnosis and therapeutic intervention contribute to reduced disease activity (*1, 2*). Though efficacious for some, current therapeutics often do not return patients to a pre-disease state, resulting in poorer quality of life and increased healthcare burden. Patients with rheumatoid arthritis (RA), a destructive systemic autoimmune disease estimated to affect 0.5-1% of the population, typically follow a course in which disease flare(s) lead to diagnosis and treatment with disease-modifying anti-rheumatic drugs (DMARDs), in a reactive response-dependent manner. Unfortunately, many patients fail treatment or relapse while on best-in-class DMARD therapies (*3*). Building on successes in type 1 diabetes therapy (*2*), an alternative proactive intervention strategy for RA targets prevention of clinical symptom onset in high risk individuals. However, this strategy is not broadly utilized because in the absence of data high-risk individuals are considered to be healthy and insufficient evidence is available to guide pre-symptom disease management.

Advances in RA risk stratification established that anti-citrullinated protein antibodies (ACPA) and rheumatoid factor (RF) may be detectable an average of 3-5 years prior to the onset of clinical RA (*4–6*), with a positive predictive value of ∼30-60% (*7, 8*). Individuals that are ACPA+ but otherwise clinically healthy are considered ‘at-risk’ for future clinical RA (at-risk individuals, ARI) and momentum has been building to better understand and modulate disease at this stage. Clinical trials in ARI have achieved some success and provide clues about important cell types and pathways accompanying transition to clinical RA. In the PRAIRI trial that enrolled ACPA+ and RF+ ARI, rituximab delayed the onset of clinical RA but did not reduce the overall rate of clinical RA development compared to placebo (*9*). In contrast, the ARIAA and APIPPRA trials demonstrated that abatacept reduced development of clinical RA within the trial periods in ACPA+ ARI (*10, 11*). Together, these trials suggest that adaptive immune responses play key roles in transition to clinical RA in ACPA+ ARI.

Cross-sectional comparisons have provided snapshots of changes in effector populations that accompany the at-risk state. Expansion of pro-inflammatory CD4 effector populations, including antigen-specific Th17 cells, along with lower glycolytic enzyme expression, are suggestive of prior activation (*12–16*). Pro-inflammatory skewing may be facilitated by elevated circulating inflammatory cytokines (*17–21*). Expanded Tph cells and ACPA specificities signify T cell-driven autoreactive B cell responses, while increased IgA+ plasmablasts and shared IgA+/IgG+ clonal families suggest a mucosal component (*13, 19, 22, 23*). There are no comprehensive, longitudinal multi-omics studies that provide a systemic understanding of the immunological changes leading to progression from at-risk to clinical RA and to highlight critical elements for diagnostics and proactive therapeutic targeting.

### ARI exhibit signs of systemic inflammation prior to the onset of clinical RA

To define molecular and cellular changes that contribute to the development of clinical RA, we studied a prospective cohort of 45 clinically-healthy ACPA+ ARI (mean follow-up of 533 days), 11 ‘early’ clinical RA (ERA) patients, and 38 ACPA- healthy controls (CON1; **Fig. 1A**, **S1A**, **tables S1-2**). During the study, 16 ARI progressed to clinical RA (‘converters’) (**fig. S1B, table S3**). Baseline (i.e. initial) ACPA levels and distribution were similar in ARI and ERA, while rheumatoid factor (RF) IgM and IgA were elevated in ERA alone (**Fig. 1B**, **S1C**). To understand the global immune state in ARI, we compared the plasma proteome from their baseline visit to CON1. We found 275 differentially abundant proteins (DAPs) in ARI (252 (92%) higher in ARI, 23 lower) (**fig. S1D, table S4**). Enrichment analysis using these DAPs identified pathways related to inflammation and cytokine, chemokine signaling (**fig. S1E, table S5**). To determine whether the baseline plasma protein alterations observed in ARI are similar to those in patients with clinical RA, we correlated the effect size changes of ARI and ERA, each relative to CON1 (**table S6**). A positive Spearman correlation between the two indicated that systemic changes in the ARI circulating proteome overlap significantly with those observed in active disease, despite absence of active arthritis in ARI (**fig. S1F**), suggesting that some molecular features of RA begin prior to clinical manifestation.

**Fig. 1:**
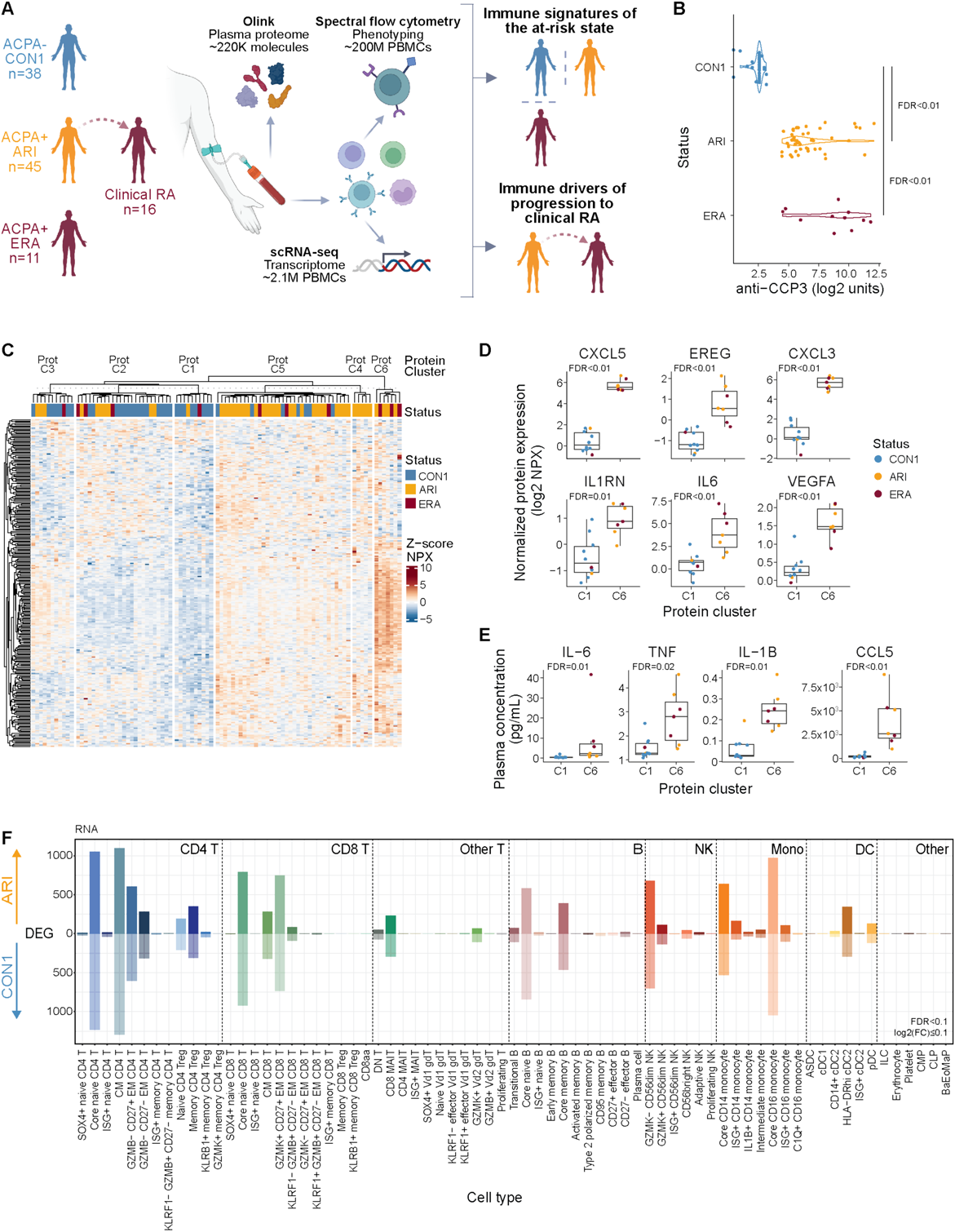
Active inflammation in ARI. (A) Overview of study and multimodal workflow. CON1, controls; ARI, at-risk individuals; ERA, early RA. (B) First sample (baseline) anti-CCP3 measurements from CON1, ARI, ERA. (C) k-means clustering (k=6) of z-scored normalized protein expression (NPX) values from differential proteins in ARI vs. CON1 (FDR < 0.1) and ERA vs. CON1 (FDR < 0.15). Rows denote proteins, columns denote baseline samples. (D) Abundance of select inflammatory plasma proteins elevated in ARI. Dots represent participant samples in each cluster. (E) Absolute concentration of select plasma proteins from participants in Prot-C1 and Prot-C6 clusters as assayed by MSD or LegendPlex. (F) Number of differentially expressed genes (DEGs; FDR < 0.1 and absolute log2 fold change ≥ 0.1) per immune cell type, elevated in ARI (above 0) or CON1 (below 0). Cell types are based on (*24*). Boxplots show median (centerline), first and third quartiles (lower and upper bound of the box) and whiskers show the 1.5x interquartile range of data. Effect sizes and *P* values for C,F were determined by linear regression models. Remaining comparisons were made using the Kruskal-Wallis test with Dunn’s post-hoc testing (B) or Wilcoxon rank-sum test (D-E). FDR values are indicated for all panels.

To evaluate heterogeneity of the identified inflammatory signature between individuals, we performed unsupervised k-means clustering of all participants (ARI, ERA, CON1) using the DAPs in ARI and ERA. This yielded 6 clusters (Prot C1-C6) in which 71.4% (40/56) of ARI or ERA segregated into Prot-C4, -C5, and -C6 (**Fig. 1C, S1G, table S7**). Clusters Prot-C6 and Prot-C1 were the most distinct in composition and exhibited the widest differences in protein abundance. Direct comparison of these two clusters uncovered additional inflammatory mediators significantly increased in ARI and ERA (**Fig. 1D, S1H, table S8**). We validated this on a subset of the identified inflammatory proteins in plasma samples from participants in clusters Prot-C1 and Prot-C6 (**Fig. 1E**). Together, these results demonstrate the presence of systemic inflammation in many ACPA+ ARI despite having no evidence of clinical RA. The observation that some ACPA+ ARI and ERA lack this proteomic signature and segregate into clusters consisting of controls highlights the heterogeneity within the ACPA+ population.

### Immune cells exhibit inflammatory gene programs in clinically healthy ARI

We hypothesized that exposure of immune cells to the inflammatory milieu in ARI would be associated with wide-ranging changes. To understand the transcriptional state of immune cells from ARI, we performed scRNA-seq on PBMCs. Three hierarchical levels of immune cell subsets were defined in the scRNA-seq data using a recently released Allen Institute for Immunology Immune Cell Atlas (*24*) and confirmed by manual review (**fig. S2A-C;** see **Methods**). We observed limited changes in relative frequencies of immune cell subsets between ARI and CON1 (**fig. S2D**, **table S9**). Despite limited changes in cell abundance, we found significant transcriptional changes in several immune cell types demonstrated by the number of differentially expressed genes (DEGs) in ARI compared to CON1 (**Fig. 1F, table S10**). The presence of numerous transcriptional changes in naive populations suggests the potential of a primed state that may contribute to disease risk, similar to that described in clinical RA (*25*). To determine how global and cell type specific transcriptional changes impact known pathways, we utilized Spectra (*26*) to analyze level 1 cell subsets (**fig. S2A**, **table S11**). We identified increased signaling downstream of inflammatory receptors specifically in CD4 T cells (**fig. S2E**) and global enrichment of glycolysis (**fig. S2F**) and oxidative phosphorylation pathways (**fig. S2G**, **table S12**). Taken together, the breadth of plasma proteomic and transcriptional changes in circulating immune cells are indicative of an ongoing inflammatory state in ACPA+ ARI.

### Progression from at-risk to clinical RA is marked by systemic immune changes and emergence of inflammatory monocyte activity

Based on previous proteomic studies (*4–6, 17–19*), we hypothesized that a significant immune triggering event would drive imminent onset of clinical RA in ARI. We focused on the 13 female ARI who progressed to clinical RA due to the availability of pre-diagnosis longitudinal samples (**Fig. 2A**). Across these samples, there was a variable range of baseline ACPA concentrations (**fig. S3A**). Twelve ARI had no appreciable longitudinal increase in ACPA levels prior to and including the time of diagnosis of clinical RA. Among clinical labs, only platelet counts increased during progression to clinical RA (**fig. S3B, table S1**). These observations prompted us to evaluate global gene expression differences. We modeled transcriptome variability over time in ARI who progressed to clinical RA relative to data from a longitudinal cohort of ACPA- healthy controls (CON2) (*24*) (**fig. S3C, table S13**). Intra-donor coefficient of variation (CV) for all detected transcripts demonstrated that 24/27 cell types had more genes with increased variability in ARI during progression to clinical RA compared to CON2 over a similar timeframe (**Fig. 2B, table S14**). In contrast, most ARI cell populations exhibited few DEGs associated with time to diagnosis (**Fig. 2C-D, table S15**). Similarly, few DAPs were associated with time to diagnosis in the plasma proteome (**table S16**). Notably, naive and central memory (CM) CD4 T cells showed the largest degree of transcriptional reprogramming, with an accompanying trend of increased abundance by flow cytometry (**Fig. 2C**, **S3D-F**, **table S17-18**).

**Fig. 2:**
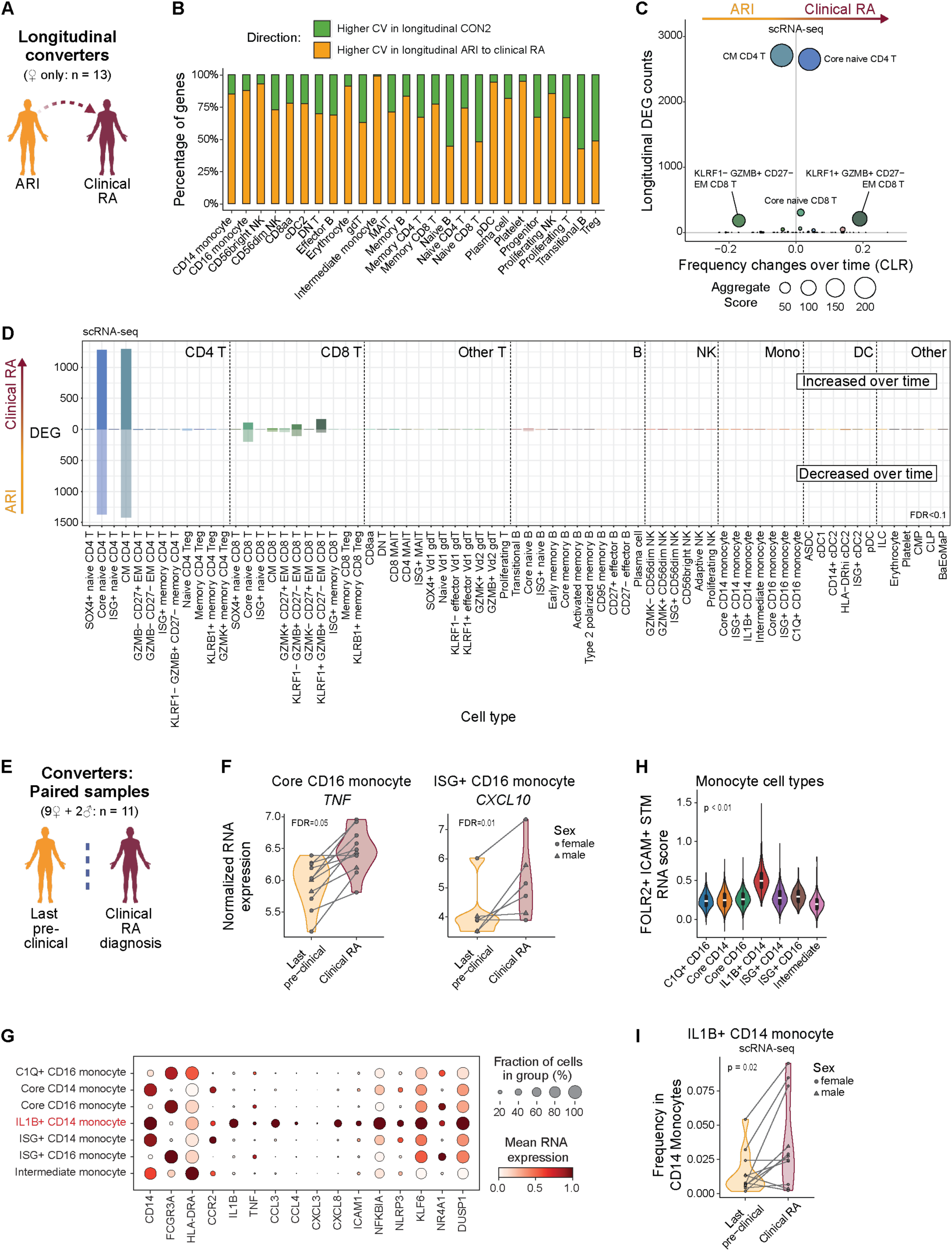
Longitudinal changes in naive and CM CD4 T cells dominate progression to clinical RA. (A) Overview of longitudinal comparison of converters, from ‘at-risk’ to clinical RA. (B) Number of genes per Allen Institute for Immunology level 2 cell type with higher average intra-donor coefficients of variation (CVs) over time in ARI who progress to clinical RA (orange) or in CON2 (green). (C) Comparison of the number of differentially expressed genes (DEGs) (y-axis) with the change in frequency over time (x-axis; centered log-ratio (CLR) transformed) as ARI progress to clinical RA. Bubble size corresponds to the aggregate score calculated by [-log(padj CLR frequency changes) x total number of DEGs]. (D) Number of DEGs from longitudinal model (FDR<0.1) per level 3 immune cell type, elevated (above 0) or diminished (below 0) in ARI progressing to clinical RA. (E-I) Overview (E) of paired comparison in converters at their last ‘at-risk’ pre-symptomatic visit vs. time of their clinical RA diagnosis. (F) Normalized RNA expression of *TNF* in Core CD16 monocytes and *CXCL10* in ISG+ CD16 monocytes. (G) Mean RNA expression of select inflammatory genes amongst monocyte level 3 cell types. (H) Gene scores calculated by comparing marker genes from FOLR2+ICAM+ RA synovial tissue macrophages (Alivernini 2020) among all monocyte cell types. (I) Frequency of IL1B+ CD14 monocytes within CD14 monocytes. Effect sizes and *P* values were determined by linear mixed effect models (C,D), paired Wald test (F), or paired Wilcoxon test (I). FDR values are indicated for all panels.

In clinical RA, cellular and transcriptional changes precede RA flares (*27*). To determine if similar changes accompany clinical RA diagnosis in converters, we compared paired samples collected at the last ‘at-risk’ pre-clinical visit (average 122 days before clinical RA diagnosis) with samples collected at the time of clinical RA diagnosis (**Fig. 2E**). Most cell types had minimal transcript changes at clinical RA diagnosis but CD16 monocytes exhibited a spike in *TNF* expression, ISG+ CD16 monocytes showed increased *CXCL10* (**Fig. 2F**), and CD95+ memory B cells showed increased *IGHA1* and *JCHAIN* expression (**fig. S3G**, **table S19**). The highest expression of *TNF* and other pro-inflammatory genes was found in IL1B+ CD14 monocytes (*28, 29*) (**Fig. 2G, S3H**) that exhibit a gene signature resembling FOLR2+ ICAM+ macrophages from RA synovial tissue (*30*) (**Fig. 2H**). *TNF* expression did not increase between visits in these cells (**fig. S3I**), but we found increased abundance of IL1B+ CD14 monocytes within total monocytes at the onset of clinical RA (**Fig. 2I, table S20**). Together, these data suggest a landscape of broad and deep changes in systemic immunity with distinct pathogenic features emerging at the time of early joint pathology.

### B cells exhibit pro-inflammatory skewing during progression to clinical RA

Both naïve and memory ARI B cells had a large number of transcriptional changes compared to CON1 B cells (**Fig. 1F**). The presence of autoantibodies and the finding of increased circulating inflammatory proteins suggested that this may reflect chronic B cell activation before onset of clinical RA. We first focused on memory B cells (MBC), particularly CD27- effector B cells often enriched in RA. Clustering ARI MBC scRNA-seq data, excluding immunoglobulin heavy (IgH) and light chain (IgL) genes, revealed expansion of a particular cluster of effector B cells (Beff-C9) during progression to clinical RA (**Fig. 3A-B, table S21**). This CD27- effector cluster was defined by genes corresponding to ‘age-associated’, Tbet+, or atypical B cells, including *ITGAX, TBX21,* and *ZEB2* (*31*), and notably shared a gene profile analogous to FCRL5+ B cells that have been associated with long-lived humoral responses (*32*) (**Fig. 3C**, **S4A**). In contrast, the other CD27- effector B cell cluster (Beff-C8) did not exhibit longitudinal changes. For comparison, a transcriptionally-related effector B cell cluster from CON2 remained stable over 2 years (**fig. S4B-C**). Beff-C9 also had a higher proportion of class-switched isotype cells (**Fig. 3D**) and higher median expression of class-switched IgH genes compared to Beff-C8 (**fig. S4D**). These data suggest CD27- atypical B cells with an activated profile expand in ARI as clinical disease approaches.

**Fig. 3:**
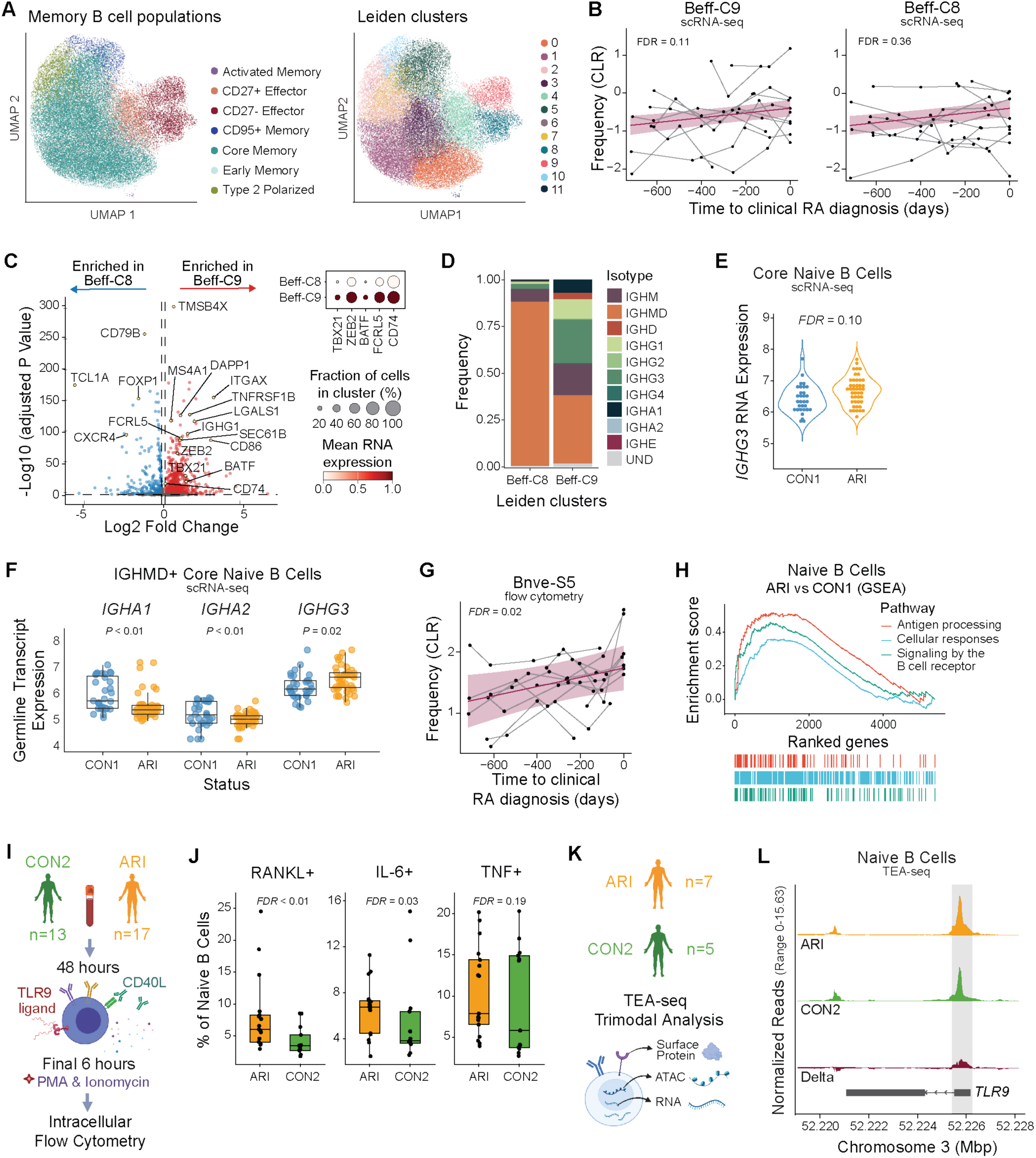
The B cell compartment exhibits a pro-inflammatory skewing during progression to clinical RA. (A) UMAP plots of memory B cells from ARI and CON1 showing B cell population labels (*left*) and Leiden clusters (*right*). (B) Centered log-ratio (CLR)-transformed frequencies of Beff-C9 (P=0.055; FDR=0.111) and Beff-C8 (P=0.359; FDR=0.359) as ARI progress to clinical RA. Each participant’s longitudinal series is connected by a gray line, with a group trendline and 95% confidence interval in purple. (C) DEGs for Beff-C8 compared to Beff-C9 in ARI samples with selected genes labeled (*left*). Dot size in heatmap (*right*) indicates the fraction of cells with positive expression for selected genes. (D) IgH isotype or undetermined (UND) identity, as frequency within each population, for Beff-C8 and Beff-C9. (E) *IGHG3* gene expression by Naive B cells (P=0.03; FDR=0.10) of ARI and CON1. (F) Normalized expression of IgH germline transcription (GLT) from IGHMD^+^ naive B cells in ARI and CON1. (G) CLR-transformed flow cytometry frequencies of Bnve-S5 as ARI progress to clinical RA, as in (B). (H) GSEA enrichment analysis with the top Reactome pathways among naive B cells of ARI compared to CON1. (I-J) B cells were stimulated *ex vivo* and analyzed by intracellular flow cytometry. Experimental workflow (I) and IL-6+, RANKL+, and TNF+ cell frequencies within the stimulated naive B cell populations of ARI and CON2 (J). (K) PBMC TEA-seq experiment overview. (L) Chromatin accessibility tracks from TEA-seq showing the *TLR9* gene in naive B cells of ARI (orange), CON2 (green) and the delta between groups (red). The gray box highlights the region containing differentially accessible peaks between groups (P=0.02; FDR=0.19). Boxplots show median (centerline), first and third quartiles (lower and upper bound of the box) and whiskers show the 1.5x interquartile range of data. *P* values were determined by a linear mixed model (B), Wald test (E,F), Wilcoxon rank-sum test (J), or zero-inflated Wilcoxon test (L). FDR values are indicated for all panels.

We hypothesized that chronic activation may influence B cell receptor (BCR) class-switching in ARI. We analyzed IgH gene expression and found higher class-switched *IGHG3* expression among ARI B cells including CD27- effector and core memory cells (**fig. S4E**). Interestingly, *IGHG3* expression differences were also observed in naive ARI B cells (**Fig. 3E**). These were confirmed to be naïve by their canonical naïve marker gene expression and by high expression of *IGHM* (**fig. S4E-G**), suggesting that elevated *IGHG3* is caused by germline transcription (GLT) due to increased transcriptional priming at that locus, despite not being class-switched. We noted that GLT was detected from all class-switched IgH genes (**fig. S5**). In IgMD+ naive cells, *IGHG3* GLT was the only IgH germline transcript showing a significant difference between ARI and CON1 (**Fig. 3F**). This suggests a greater likelihood that naïve B cells will class-switch to the IgG3+ isotype upon activation, an intriguing finding given that patients with RA and other select autoimmune diseases are reported to have higher total IgG1 and IgG3 levels (*33*). Detailed evaluation of B cells by flow cytometry revealed a subset of naïve B cells (Bnve-S5) that expanded during progression to clinical RA in ARI (**Fig. 3G**, **S4H**), a feature previously reported to correlate with reduced PAX5 expression (*13*). Indeed, *PAX5* expression within naive ARI B cells was reduced (**fig. S4I**). A broader evaluation of the transcriptome in naive ARI B cells showed enrichment for transcripts associated with cellular responses to external stimuli, antigen processing and presentation, and BCR signaling when compared to CON1 (**Fig. 3H**).

To determine whether the broadly activated molecular profile of naive ARI B cells represents a functionally “primed” state, we stimulated ARI and CON2 B cells with CPG and MEGACD40L to stimulate TLR9 and CD40, mimicking microbe-response signals and T cell help. We evaluated cytokine expression by B cell subsets using intracellular flow cytometry (**Fig. 3I**). Naive B cells from ARI had higher frequencies of IL-6+, RANKL+, and TNF+ cells compared to controls (CON2) after stimulation (**Fig. 3J**) suggesting they are functionally primed. Using TEA-seq, a tri-modal single-cell assay combining CITE-seq (surface proteins and scRNA-seq) and ATAC-seq (*34*) (**Fig. 3K**), we evaluated resting naive B cells for expression of genes potentially associated with our observed *in vitro* activation differences. This showed greater accessibility at the *TLR9* promoter in ARI vs. CON2 naïve B cells (**Fig. 3L**). This suggested potential transcriptional changes affecting *TLR9* but we were unable to confirm this due to transcripts for most TLRs, including TLR9, not being well captured in the scRNA-seq data. Collectively, naive B cells of ARI may have enhanced BCR signaling and antigen presentation, and are primed for IgG3 class-switching and elevated proinflammatory cytokine and RANKL secretion. Together these data show broad activation across naïve and memory ARI B cell populations with transcriptomic and functional evidence demonstrating that this leads to naïve B cells being in a primed state.

### Tfh17 CD4 T cells expand during progression to clinical RA

Given the role of CD4 T cells in B cell activation and class switching, we next evaluated ARI effector T cell populations. In fact, CM CD4 T cells had the highest number of DEGs in converters as they progressed to clinical RA (**Fig. 2D**). Analyzing the transcriptome at the population level by pseudobulk analysis, we found a signature of increased T cell activation, including downregulation of *CD3G* and *CD247* (CD3Z) and upregulation of genes related to cytokine and antigen receptor signaling (*STAT5B*, *STAT2*, *STIM2*, *AKT3*, *CD28* and *FOSB*) as ARI approached disease diagnosis (**Fig. 4A-B, S6A, table S22**). This was accompanied by a trend of increasing abundance of a CD4 memory T cell subcluster by flow cytometry (**fig. S3F**, **table S17-18**), indicating that T cells are activated during disease development.

**Fig. 4:**
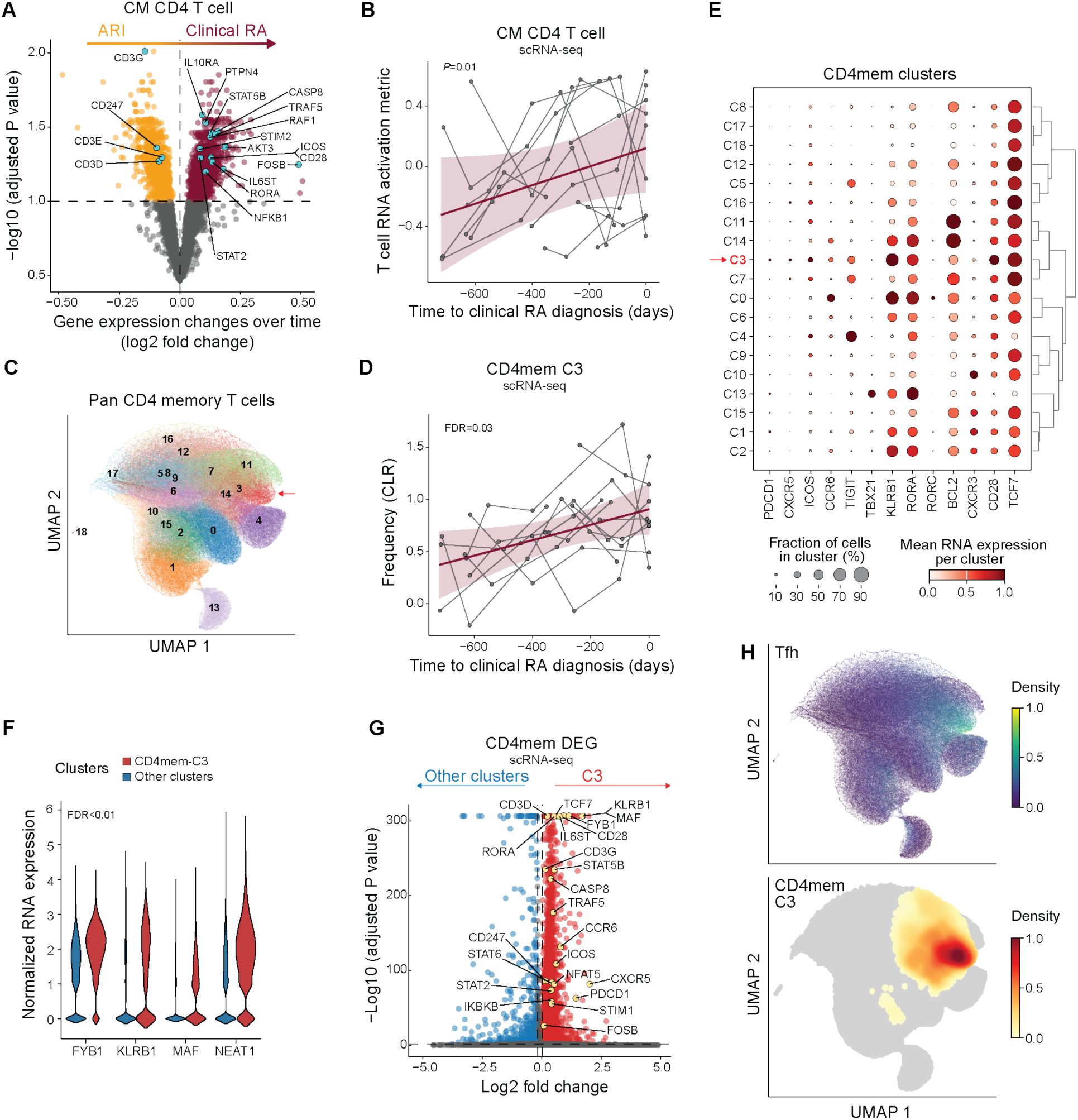
Expansion of effector and memory T cells with pathogenic signatures during progression to clinical RA. (A) RNA expression differences in central memory (CM) CD4 T cells over time in ARI (orange) who progress to clinical RA (purple). Genes associated with T cell activation are noted. (B) T cell RNA activation metric in CD4 CM over time as ARI progress to clinical RA. Each participant’s longitudinal series is connected by a gray line, with a group trendline and 95% confidence interval in purple. (C) UMAP showing Leiden clustering of non-negative matrix factorization (NMF)-projected CD4 reference gene weights on pan CD4 memory T cells (CD4mem). Cluster 3 (CD4mem-C3) is indicated (arrow). (D) Frequency over time in CD4mem-C3 cells as ARI progress to clinical RA, as in (B). (E) Mean RNA expression of select genes across CD4mem clusters. (F) Normalized RNA expression of genes that promote differentiation to Tfh and Th17 cells in CD4mem-C3 (red) vs. remaining CD4mem clusters (blue). (G) Differentially expressed genes between CD4mem-C3 (red) and remaining CD4mem clusters (blue). Select genes associated with T cell activation are labeled. (H) Cells expressing Tfh gene program are distinguished based on the NMF projection using a pre-computed weight matrix of CD4 T cell population from Yasumizu et al. 2024. For comparison, a UMAP density plot of cluster CD4mem-C3 is shown below. *P* values were determined by linear mixed models (A, B, D) or the Wilcoxon rank-sum test (F-G). Nominal *P* value is indicated for (B). FDR values are indicated for (A, D, F, G).

To understand how the activation signature impacts effector phenotypes during clinical RA development, we projected our CD4 T cell scRNA-seq data from converters onto a human CD4 gene program reference (*35*) (**Fig. 4C**) and analyzed memory clusters for abundance and DEGs. Cluster 3 (CD4mem-C3) displayed significantly increased abundance during progression to clinical RA (**Fig. 4D**, **table S23**). Clusters CD4mem-C7, -C11 and -C14 also trended towards increased abundance (**fig. S6B**). Gene expression suggested CD4mem-C3 contained a subset of Tfh cells that share a Th17 gene signature (i.e. Tfh17), as evidenced by *PDCD1*, *CXCR5*, and CD28, along with *CCR6*, *KLRB1*, and *RORC* expression (**Fig. 4E, table S24**), but are distinct from Th17 cells (CD4mem-C0). CD4mem-C3 was also the predominant cluster expressing genes important for Tfh and Th17 differentiation and functionality (*36–38*) and general TCR signaling (*39*) (**Fig. 4F**, **S6C**). Eighty percent of the CD4mem-C3 population labeled as CM CD4 T cells (**fig. S7A**), overlapping in UMAP space and having similar DEGs (**Fig. 4A, 4G, S7B**). Further, CD4mem-C3 aligned in UMAP space to the Tfh gene program from a reference dataset (*35*) (**Fig. 4H**, **S7C-D**). These finding suggest that the observed increase of CM CD4 T cell abundance as ARI approach clinical RA is driven predominantly by changes in Tfh/Tph cells that share features of Th17 cells (Tfh17). Paired with pre-activated primed B cells, these may contribute to autoantibody development in RA.

### Naive T cells also exhibit activation signatures during progression to clinical RA

Similar to the B cell compartment exhibiting activation of both effector and naïve populations, ARI T cells demonstrate a similar pattern (**Fig. 1F**). In converters, naïve CD4 T cells are one of only two populations that show a large number of DEGs accompanying progression to clinical RA, suggesting ongoing T cell proliferation and/or activation. Indeed, in naive T cells, we detected downregulation of CD3 complex transcripts and upregulation of cytokine and antigen receptor signaling components (**Fig. 5A-D**), the latter including STATs, *STIM2*, and *CD28* (**fig. S8A-B**). Gene set enrichment analysis (GSEA) indicated activation pathways, including NFAT and TGFβ signaling, in both CD4 and CD8 naïve T cells (**fig. S8C-D, table S25**). Together, these results are indicative of activation and suggest that naive T cells from ARI may be continually subjected to immunomodulatory stimuli during progression to clinical RA.

**Fig. 5:**
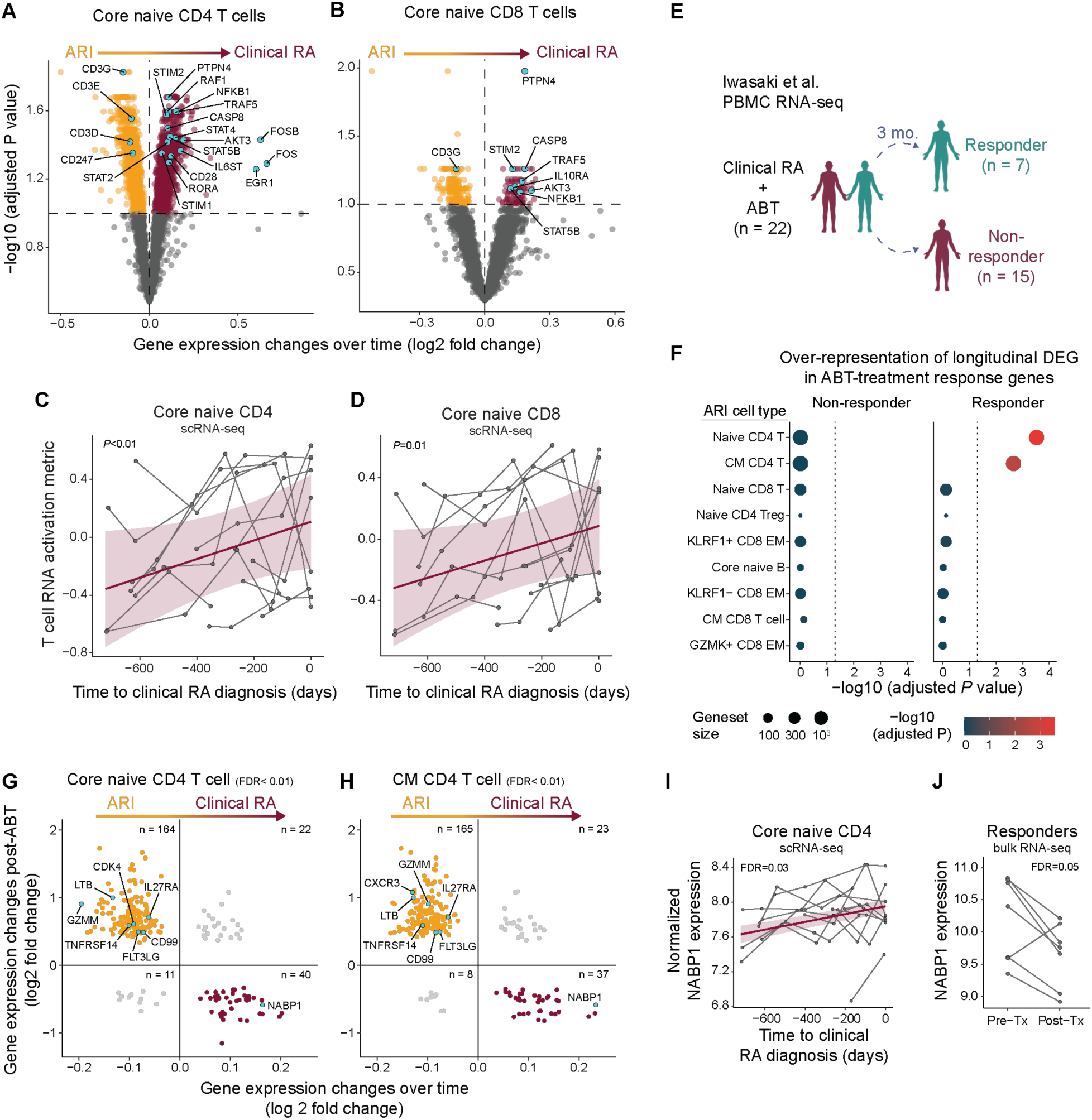
Activation signature in naive T cells during progression to clinical RA. (A-B) RNA expression differences in core naive CD4 (A) or CD8 (B) T cells over time in ARI (orange) who progress to clinical RA (purple). Genes associated with T cell activation are annotated. (C-D) T cell RNA activation metric in core naive CD4 (C) or CD8 (D) T cells over time as ARI progress to clinical RA. (E-H) Longitudinal DEGs as ARI progress to clinical RA were assessed within the context of RA patients with efficacious (responders) or non-efficacious (non-responders) clinical response to abatacept (ABT) treatment (from Iwasaki et al. 2024). (E) Overview of the analysis strategy. (F) Over-representation of ARI cell type-specific longitudinal DEG amongst ABT-treatment response DEG. (G-H) Significant DEGs in ABT responders compared to DEG changes over time as ARI progress to clinical RA in core naive CD4 T cells (G) and CM CD4 T cells (H). Genes (dots) previously implicated in RA-like disease are labeled. (I-J) Normalized RNA expression of NABP1 over time as ARI progress to clinical RA (I) and pre- vs. post-ABT therapy in RA patients (J). *P* values were determined by linear mixed models (A-D, I), hypergeometric enrichment tests (F), McNemar’s Chi-squared test (G-H), Wald test (J). Nominal *P* values are indicated for (C-D). FDR values are indicated for (A, B, F-J).

To understand the relevance of T cell changes observed in converters, we compared their transcriptomic signatures to a whole blood transcriptome dataset generated in a clinical trial of abatacept (ABT; CTLA4-Ig) in patients with active RA (*40*) (**Fig. 5E**). ABT is hypothesized to dampen T cell costimulation in RA (*41*). Notably, naive and CM CD4 T cell gene signatures found in converters over time were significantly enriched specifically in ABT responders prior to treatment (**Fig. 5F, fig. S8E-F**). Indeed, the majority of genes in the naive and CM CD4 T cell gene signatures were inversely correlated between ARI who progress to clinical RA and RA responders post-ABT (**Fig. 5G-H**), suggesting that mechanisms relevant to clinical RA contribute to disease pathogenesis in ARI, and that these can be reversed by a therapy targeting T cell activation. We highlight ABT-driven changes in genes previously implicated in RA-like disease (*CD8a*, *CD99*, *CDK4*, *CXCR3*, *FLT3LG*, *GZMM*, *LTB*, *TNFRSF14*) and those related to the Th17 pathway (*IL27RA*, *NABP1*) (**Fig. 5G-J**, **table S26**). These results support the hypothesis that T cell activation is a critical element in progression of ARI to clinical RA, and provide mechanistic evidence supporting the role of ABT in delaying onset of clinical RA (*10, 11*).

### Epigenetic changes in naive CD4 T cells are linked to NFAT-calcium activation and Tfh bias in ARI

Given the systemic inflammation and longitudinal increase of Tfh17-like effector cells in ARI, we hypothesized that naive CD4 T cells are in a state of heightened activation and are epigenetically poised to preferentially differentiate into pathogenic effector cells. We performed TEA-seq on PBMCs from a subset of ARI and matched CON2 participants (**Fig. 3K**, **table S1**), labeled major immune cell types (**fig. S9A;** see **Supplementary methods**), then integrated the plasma proteome, transcriptome, surface proteins, and TF activities (inferred by motif accessibility) from naive CD4 T cells using multi-omics factor analysis (MOFA) (*42*). Factor 1 explained the highest variation in the transcriptome (**Fig. 6A**) and differentiated ARI from CON2 (**Fig. 6B)**, with transcripts encoding calcium responsive proteins (*NFAT5*, *NFATC3*, *NFATC2*) and signaling proteins (*PPP3CC*, *PPP3CA*, *STIM1*, *STIM2*) elevated in ARI (**Fig. 6C**). Moreover, within Factor 1, we detected increased NFAT TF motifs and decreased FOX TF motifs in accessible chromatin regions of ARI (**Fig. 6D, S9B**). These results confirm our earlier observation that naive CD4 T cells from ARI are in an activated state.

**Fig. 6:**
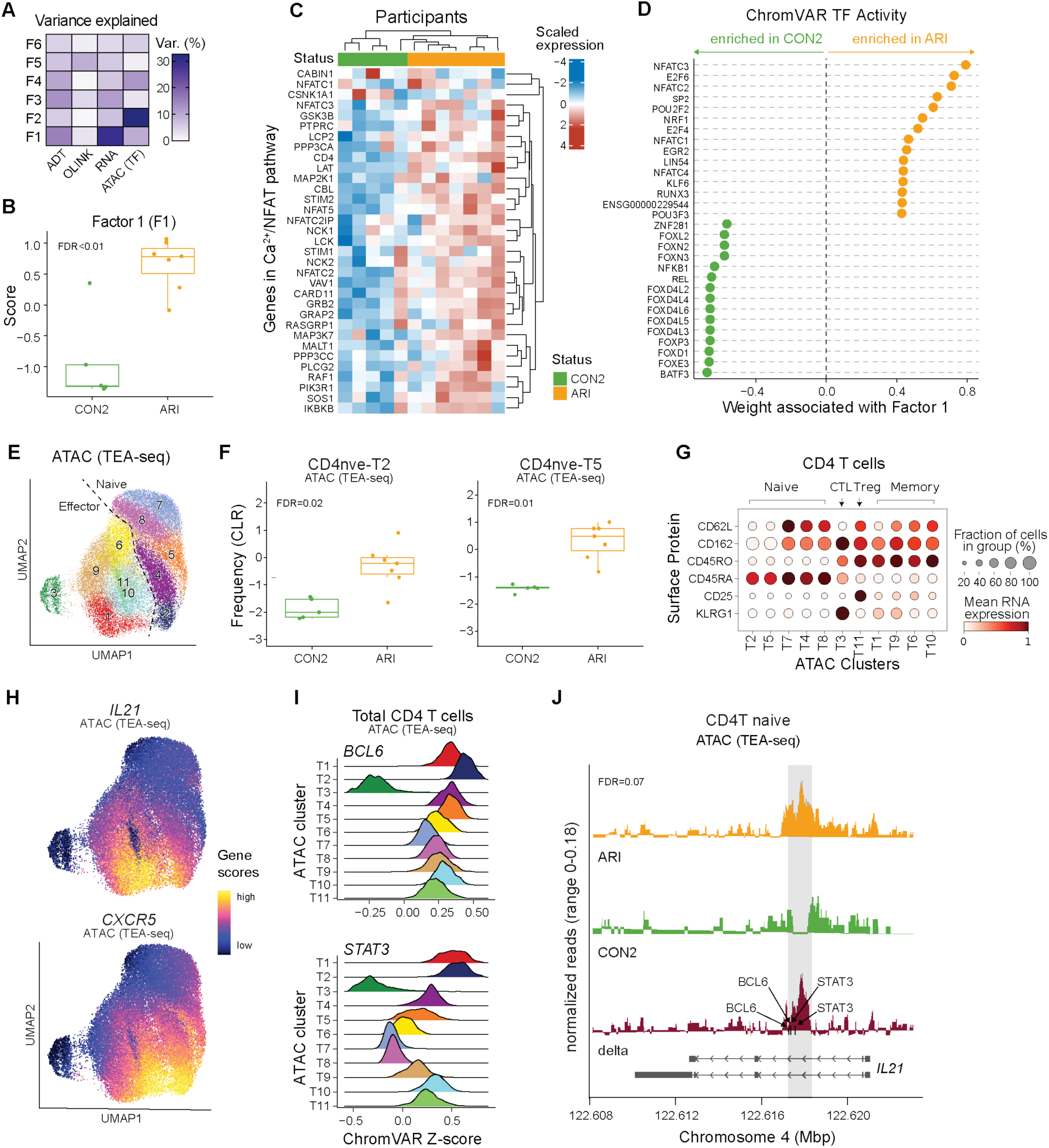
Epigenetic changes in naive CD4 T cells support activation and Tfh bias in ARI. PBMC TEA-seq experiment in a subset of ARI and CON2 samples was performed as in Fig. 3K. (A) Percentage of variance in each modality (surface protein, plasma protein, RNA, ATAC) explained by Multi-Omics Factor Analysis (MOFA) factors. (B) Factor 1 scores between ARI and CON2. (C) Scaled normalized expression of select genes in Calcium–Calcineurin–NFAT pathway in ARI and CON2. (D) Inferred accessibility for the top 15 transcription factors (TFs) positively or negatively associated with factor 1, ranked by weight. (E) Louvain clusters in CD4 T cells by ATAC modality in TEA-seq. (F) Centered log-ratio (CLR)-transformed frequencies of ATAC clusters CD4nve-T2 and CD4nve-T5 in CD4 T cells. (G) Mean surface protein expression of select markers differentiating CD4 naive, memory, Treg, and cytotoxic CD4 T cells (CTL) across ATAC clusters. (H) ATAC UMAP overlaid with inferred gene activity scores calculated by ArchR for *CXCR5* and *IL21*. (I) ChromVAR TF activity z scores of *BCL6* and *STAT3* in CD4 T cells. (J) ATAC signal in ARI (orange), CON2 (green), and delta (red) at the *IL21* locus. The gray box highlights a 500bp region containing differentially accessible peaks between ARI and CON2 (chr4: 122,617,500-122,617,999). Black arrows indicate the motif locations of BCL6 and STAT3 binding sites. Gene bodies are displayed on the bottom. Boxplots show median (centerline), first and third quartiles (lower and upper bound of the box) and whiskers show the 1.5x interquartile range of data. *P* values were determined by linear models (B, F) or zero-inflated Wilcoxon test (J). FDR values are indicated.

We next compared accessible regions in naive CD4 T cells between ARI and CON2, and identified 3,159 differentially accessible peaks, including 2,200 in promoter regions (**table S27**). Louvain clustering on all CD4 T cells based on the ATAC modality alone (**Fig. 6E**) revealed two CD45RA+ naive clusters (CD4nve-T2, -T5) with a higher frequency in ARI (**Fig. 6F**) and three CD45RO+ memory clusters (CD4eff-T6, -T9, -T10) with a lower frequency (**fig. S9C**). CD4nve-T2 and CD4nve-T5 were distinct in the epigenetic space (**Fig. 6E**), though not in the RNA or ADT space (**fig. S9D**). Clusters CD4nve-T2 and CD4nve-T5 exhibited effector-like phenotypes of lower CD62L and CD162 surface protein expression (**Fig. 6G**), and high chromatin accessibility at the *CXCR5* and *IL21* loci compared to other naive clusters (**Fig. 6H**). Notably, CD4nve-T2, which had evidence for an activated or pre-activated state shared with CD4eff-T1 as indicated by TF motif enrichment for SMADs, STATs, and AP1 (**fig. S9E, table S28**), had the highest ChromVAR scores for BCL6 and STAT3 predicted activity (**Fig. 6I**). Moreover, although minimal *IL21* transcript was detected (**fig. S9F**), an accessible 500bp intronic region in the *IL21* locus was present only in naive CD4 T cells from ARI (**Fig. 6J**). This region was shown to be more accessible in human tonsil Tfh cells (*43*) and overlaps an ENCODE putative enhancer-like structure that contains motifs for key TFs that drive Tfh, Tph differentiation, including BCL6 and STAT3 (*44*) (**Fig. 6I**). These results identify a putative regulatory region in naive CD4 T cells that is permissive to induction of IL-21 specifically in ARI, and links naive T cell activation in ARI to Tfh/Tph cell differentiation.

## Discussion

Development of ACPAs are strongly predictive for development of future clinical RA (*4, 5*), but immune changes underlying progression from the ACPA+ at-risk state to clinical disease remain unclear. Here, we significantly expand prior knowledge of immune alterations that precede clinical RA, by demonstrating extensive inflammatory changes in ARI using systems immunology applied to a longitudinal, prospective at-risk cohort. These proinflammatory features mirror alterations we observed in ERA and those reported in active arthritis, including in inflamed synovial tissue (*28–30*). Together, these results support the concept that disease begins earlier than clinically appreciated, with ACPA+ ARI having many systemic features of active RA-like inflammation preceding development of clinical symptoms, with broad implications for understanding autoimmune pathogenesis.

As a hallmark of seropositive RA, ACPAs are one of the most extensively profiled features of autoimmunity. Nonetheless, we have an incomplete understanding of the role of B cells in the development of clinical RA. Prior studies show that autoreactive and activated B cell populations contribute to clinical RA symptomology and progression (*45, 46*), partially attributable to their production of autoantibodies and proinflammatory factors. We identified a subset of CD27- ITGAX+ effector memory B cells that increase in abundance in ARI during progression to clinical RA. This memory subset showed transcriptional features suggestive of chronic BCR engagement, including elevated class-switching, and a gene program associated with long-lived humoral immunity after vaccination (*32*). In clinical RA, subsets of CD11c+ (encoded by *ITGAX*) atypical memory B cells are clonally related to expanded plasma cells in the periphery and synovial tissue (*47, 48*), highlighting this population as a potentially interesting target in preventative therapies.

Notably, we found that naive B cells in ARI also showed evidence of activation, previously recognized as a harbinger of disease flares in active RA (*27*). In our study a subset of naive B cells increased in abundance during disease progression and total naive cells were primed for IgG3 class-switching. In addition, greater proinflammatory cytokine and RANKL secretion was observed following *in vitro* stimulation of naive B cells, expanding on literature showing activated memory B cells in active RA produce more RANKL than controls and promote osteoclastogenesis *in vitro* in a RANKL-dependent manner (*49*). These data suggest that not only the production of ACPA but ongoing B cell activation is a feature of individuals progressing to clinical RA.

Naive T cells also exhibit constitutive activation in clinically recognizable RA (*25*). We note increased activation signatures in naive CD4 T cells from ARI, which we hypothesize result from the chronic inflammatory milieu of ARI. In memory T cells we found expansion of a subset that has the composite transcriptional features of both Tfh/Tph and Th17 helper T cell subsets similar to Tfh17 cells, an IL-17A-producing subset of Tfh. Tfh17 cells efficiently promote B cell function and are correlated with circulating autoantibodies in juvenile dermatomyositis and Hashimoto’s thyroiditis (*50, 51*). The Tfh17 cells we identified in ARI expressed MAF, a key transcription factor promoting Tfh and Th17 cell fate, including expression of CXCR5 and IL-21 (*36, 52, 53*). These data expand upon literature implicating Tfh, Tph in the pathogenesis of RA (*54*).

Using a published dataset, we show that the gene signature present in naive and CM CD4 T cells in ARI that convert to clinical RA was elevated before treatment in active RA patients that subsequently responded to abatacept and decreased after 3 months on therapy (*40*). Notably, the Tfh17 cluster that increased in ARI during progression to clinical RA also expressed the highest level of CD28, a subset of Tfh proposed to be a target of CTLA-4 modulation (*55, 56*). These data suggest that the inflammatory T cell signatures we identified may be contributing to disease pathogenesis in ARI. Though causative mechanisms can’t be tested with these data, these results warrant further investigation.

We sought to understand mechanisms that might link perturbation in naive T cells with changes noted in effector T cells. Trimodal profiling of cell surface proteins, transcriptome, and epigenome provided the opportunity to better understand molecular mechanisms underlying naive T cell activation and Tfh17 expansion in ARI. Pre-existing transcriptional and epigenetic landscapes in naive T cells can skew their effector potential towards Tfh development (*57*). Consistent with previous reports in clinical RA (*25*), naive CD4 T cells in ARI showed activation through calcium and TCR signaling. Epigenetically, this was associated with a subcluster of cells that demonstrated effector-like chromatin accessibility profiles, including more accessible BCL6 and STAT3 motifs, known drivers of Tfh differentiation (*58*), as well as a more accessible regulatory region at *IL21*. These results suggest a mechanism whereby Tfh17 expansion in ARI may result from a pre-existing bias in naive cells, highlighting the possibility that chronic inflammation in ARI is a feed-forward process that culminates in joint pathology. Furthermore, the signature of heightened T cell activation, Tfh expansion and priming for IL-21 production, captured in the periphery, may contribute directly to elevated B cell activation detected in ARI.

Despite a plethora of changes in ARI relative to healthy controls and in ARI over time as they progress to clinical disease, we observed limited changes in circulating immune cells at the time of RA diagnosis compared to the prior visit (∼122 days), despite their emerging clinical symptoms. At clinical RA onset, TNF expression in CD16 monocytes spiked while a population of CD14 monocytes acquired a TNF+IL1B+ phenotype. The IL1B+ monocyte population resembled FOLR2+ ICAM+ synovial tissue macrophages (*30*). Whether the circulating TNF-expressing monocytes we identified traffic to the joint prior to clinical symptoms or as a consequence of existing joint inflammation remains to be determined and could be informative for understanding the initiation of joint pathology.

One limitation of this study is participants were predominantly white, non-Hispanic individuals, so further studies would be required to generalize the findings to other demographic populations. Analysis of paired synovium was not performed as part of this study. We did, however, identify signatures of immune populations in ARI PBMC that overlapped those reported in cells isolated from active arthritis synovium (*29, 30*). We modeled longitudinal changes during progression of ARI to clinical RA by comparing intra-donor data from pre-symptomatic timepoints through clinical disease onset, rather than performing inter-donor comparisons with ARI that have not developed clinical RA. The latter group is inherently heterogeneous and lacks a shared timeline in disease status, making direct comparisons difficult.

While the underlying drivers of the observed immune activation are not clear, several lines of evidence suggest that disruption of mucosal surfaces may lead to increased transit of pathogen derived products promoting innate immune cell activation and priming of adaptive cells (*59–62*). This will require future studies. Whether the inflammatory process precedes the development of autoantibodies and promotes the break in tolerance, or the autoantibodies develop first, subsequently driving an inflammatory response, remains an important topic of future investigation. Taken together, these data provide an unmatched, systems view of the immunologic features of ACPA+ ARI and provide numerous potential biomarkers that may be used to predict which ARI will develop clinical RA. Given that clinical trials for RA prevention are progressing in ARI (*9–11, 63*), these findings also suggest additional opportunities and strategies for prevention, potentially focusing on interrupting T-B cell interactions, Tfh/Tph or Th17 function, and/or cytokine inhibition, such as blocking IL21.

## Materials and Methods

### Study Design

#### Overview of study participants

The following groups were included in this study: (1) At-Risk Individuals (ARI), defined as having serum ACPA positivity using the anti-cyclic citrullinated peptide-3 (anti-CCP3 IgG ELISA, Werfen); (2) Early RA (ERA), defined as having on physical examination ≥1 swollen joint consistent with synovitis and anti-CCP3 positivity, with their initial study visit taking place within 1 year of the initial confirmation of synovitis by a rheumatologist; (3) ACPA- Controls (CON1), defined as being anti- CCP3 negative and recruited at the University of California San Diego and the University of Colorado; (4) Longitudinal healthy ACPA- controls (CON2), defined as being anti-CCP3 negative and recruited at the Benaroya Research Institute (*24*). Clinical data associated with the study were finalized as of December 18, 2023.

#### Group recruitment and enrollment

ARI, ERA and CON1 participants were identified and recruited by the University of California San Diego (UCSD) and the University of Colorado Anschutz Medical Campus (CU) as part of the Allen Institute for Immunology-UCSD-CU Transition to Rheumatoid Arthritis Project (ALTRA) Project. ARI were identified through anti-CCP3 testing of first-degree relatives of individuals with clinical RA, health-fair testing, research outreach to campuses and communities, and through evaluation of rheumatology clinic referrals. All ARI were positive at enrollment for the anti-CCP3 test (20 units) and had no history of or examination evidence of synovitis at the time of enrollment as determined by a rheumatologist or trained study nurse’s physical examination of 68 joints (joint tenderness in absence of swelling was not an exclusion). Additional exclusions for the ARI included use of immunomodulatory therapy other than short courses of corticosteroids for non-inflammatory arthritis conditions (e.g. asthma, although use of corticosteroids for gout was allowed). ERA were recruited from rheumatology clinics at UCSD and CU and included individuals with ≥1 swollen joint consistent with synovitis identified by a rheumatologist, and their ALTRA study visit occurred within 1 year of their initial diagnosis of clinical RA by a rheumatologist. Furthermore, of the enrolled Early RA participants 11/11 (100%) met the 2010 American College of Rheumatology/European Alliance for Associations of Rheumatology (ACR/EULAR) RA classification criteria. Recruitment of both ARI and ERA was facilitated through use of signage, flyers and social media campaigns (*64*). CON1 were identified as being anti-CCP3 negative through testing of first-degree relatives of patients with RA as well as individuals at health-fairs, campus and community outreach and rheumatology clinics which are similar venues where the anti-CCP3 positive individuals were identified. Exclusion criteria for these participants included inflammatory arthritis and/or the use of immunomodulatory agents.

CON2 participants were recruited from the Seattle area by the Benaroya Research Institute as part of the Sound Life Project and sampled longitudinally (*24*). Individuals were excluded if they had a history of diagnosed chronic disease, autoimmune disease, severe allergy, or chronic infection. All participants had anti-CCP3 levels < 20 units.

#### Ethical considerations

All studies were approved by ethical review boards at UCSD, CU, Benaroya Research Institute and the Allen Institute for Immunology. All participants provided written informed consent prior to participation in these studies.

#### Sample collection

Consistency in sample collection and handling between the three sites was through the use of a common lab manual, common protocols, on-site training prior to the start of work, and regular coordination meetings. All samples were assayed at the Allen Institute, except for plasma proteomics, which was outsourced (Olink).

Blood was drawn into BD sodium heparin (NaHeparin) vacutainer tubes for PBMC or K2-EDTA vacutainer tubes for plasma. PBMC isolation and plasma processing were started within 2 hours post blood draw. For PBMC isolation, upon arrival at the processing lab all NaHeparin tubes for each donor were pooled into a sterile polystyrene pooling receptacle, gently swirled ∼30 times until fully mixed, and combined with an equivalent volume of room temperature PBS (ThermoFisher). PBMCs were isolated using Leucosep tubes (Greiner Bio-One) loaded with 15 mL of Ficoll-Paque Premium (GE Healthcare) to which a 3 mL cushion of PBS had been slowly added on top of the Leucosep porous barrier. 24–30 mL diluted whole blood was slowly added to the tube and spun at 1000 x g for 10 min at 20°C with no brake. PBMCs were recovered by quickly pouring all supernatant volume above the barrier into a sterile 50 mL conical tube; 15 mL cold PBS+0.2% BSA (Sigma; “PBS+BSA”) was added, and the cells were pelleted at 400 x g for 5–10 min at 4–10°C. The supernatant was quickly decanted, the pellet dispersed by flicking the tube, and the cells washed with 25–50 mL cold PBS+BSA. Cell pellets were combined, if applicable, and the cells were pelleted as before. PBMCs were resuspended in 1 mL cold PBS+BSA per 15 mL whole blood processed and counted with a Cellometer Spectrum (Nexcelom) using Acridine Orange/Propidium Iodide solution. PBMCs were cryopreserved in 90% FBS (ThermoFisher) / 10% DMSO (Fisher Scientific) at a target of 5 x 10^6^ cells/mL by slow freezing in a Coolcell LX (VWR) overnight in a -80°C freezer followed by transfer to liquid nitrogen.

### Assays

#### Autoantibody and inflammatory marker testing

Anti-citrullinated protein antibodies (ACPA) were tested using the anti-cyclic citrullinated peptide antibody-3 (anti-CCP3 IgG) assay using an ELISA platform (QuantaLite, Werfen) with a cut-off for positivity based on the manufacturer’s suggestion of ≥20 units. Rheumatoid factor (RF) immunoglobulin (Ig) A and IgM were tested using an ELISA platform (QuantaLite, Werfen) with cut-offs for positivity based on the clinical laboratory’s internal cut-offs set using 80 controls (>1.7 International Units for RFIgA and >6.7 International Units for RFIgM). High sensitivity C-reactive protein (hsCRP) testing was performed using nephelometry with results in milligrams per liter (mg/L). Erythrocyte sedimentation rates (ESR) were performed using clinical lab methodologies with results in millimeters per hour (mm/hr). Anti-CCP3, RFIgA and RFIgM tests were performed on all ALTRA participants (ARI, ERA, CON1) as well as CON2, and all testing for these autoantibodies was performed at the University of Colorado in the Exsera Biolabs, a College of American Pathologists and Clinical Laboratory Improvement Amendments (CAP/CLIA) certified laboratory. hsCRP and ESR testing were performed using local clinical laboratories at UCSD, CU and BRI.

#### Shared epitope testing

For ALTRA participants, presence of the shared epitope was determined by high resolution polymerase chain reaction (PCR) at the University of Colorado ClinImmune Laboratory. Alleles considered to contain the shared epitope include: HLA-DRB1 *01:01, *01:02, *04:01, *04:04, *04:05, *04:08, *04:09, *04:10, *04:13 and *10:00.

#### Plasma proteomics

For plasma isolation, the K2-EDTA source tube was gently inverted 10 times, centrifuged at 2000 x g for 15 min at 20°C with a brake of 1, and 80–90% of the plasma supernatant was removed for immediate freezing at −80°C. Plasma was assayed after the first freeze/thaw. Plasma samples were assayed on the Olink Explore 1536 platform, which uses paired antibody proximity extension assays (PEA) and next-generation sequencing to measure the relative expression of 1,472 total protein analytes (1463 unique) per sample. Samples were randomized across plates to achieve a balanced distribution of age and sex. Longitudinal samples from the same participant were run on the same plate. Plasma bridging controls (12) were included in each of 7 batches and used for cross-batch normalization.

#### Mesoscale Discovery (MSD) and LegendPlex Targeted Assays

Plasma concentrations of IL-6 and TNF were measured using the V-PLEX Human Proinflammatory Panel 1 kit (MSD #K15049D-2), IL-1B using the S-PLEX Human IL-1B kit (MSD #K151ADSS), and CCL5 using the LEGENDPplex Human Proinflammatory Chemokine 1-plex kit (BioLegend #741083) according to the manufacturer’s protocol. For MSD assays, 25µl of undiluted plasma samples were utilized and plates were read on a Meso Sector S 600MM instrument. Sample raw data and standard curves were derived using Discovery Workbench v4.0.13. For LEGENDplex assays, 25µl of diluted plasma (1:50 in assay buffer), were utilized and samples were analyzed on a Cytek Aurora flow cytometer. Data acquisition and analysis were performed according to BioLegend’s instrument setup (for Cytek Aurora) (https://www.biolegend.com/Files/Images/BioLegend/legendplex/instructions/Setup_Procedure_for_Cytek_Aurora_and_Northern_Lights_2_Lasers.pdf) and analysis software suite (https://www.biolegend.com/en-us/immunoassays/legendplex/support/software).

#### Flow cytometry and scRNA-seq

For prospective flow cytometry and scRNA-seq, PBMCs were assayed after the first thaw according to a previously published method (*65*). In most cases, flow cytometry and scRNA-seq data were collected from the same vial of cells. For flow cytometry, cells were labeled with a selection of antibodies targeting 56 surface proteins organized into 4 panels and data were collected on a Cytek Aurora, as previously described (*66*). For scRNA-seq, samples were hashed and libraries generated using a modified 10x Genomics assay, as previously described (*65*). Hashed data processing was carried out using BarWare (*67*) to generate sample-specific output files.

#### TEA-seq

Seven ARI and six age-matched female CON2 donors were selected (**table S1**). One CON2 donor, who tested positive for anti-CCP3 (>20 units), was excluded from further analysis. TEA-seq library preparation was performed as described previously (*34, 68*). In brief, all samples were thawed and stained with the sample specific cell hashing antibodies then processed in the same batch. Cells were sorted to remove dead cells, debris, and neutrophils. A panel of 166 target-specific barcoded oligonucleotide-conjugated antibodies (ADT) (BioLegend TotalSeq™-A Human Universal Cocktail, V1.0 and 3 additional antibodies) were used in the present experiment (**table S29**). Individual ATAC, RNA, hashtag oligonucleotide (HTO) and ADT libraries were prepared, sequenced and processed as described previously (*34*).

### Intracellular Flow Cytometry

#### Cell Assay, Processing and Antibody Staining

Total PBMCs were suspended in RPMI 1640 supplemented with 20% heat-inactivated fetal bovine serum (FBS), 1% penicillin-streptomycin, L-glutamine, HEPES, sodium pyruvate, glucose and 2-Mercaptoethanol (Gibco; 50 μM) at 4 × 10^6^ cells/mL and rested for 30 minutes. For each subject’s sample, half of the PBMC suspension was then stimulated for 48 hours with CpG ODN 2006 (Invivogen; 10 μg/mL) and MEGACD40L (Enzo; 100 ng/mL) or the other half was cultured without stimulation in a sterile 96-well U-bottom culture plate (Falcon) at 37°C and 5% CO2. Phorbol 12-myristate 13-acetate (PMA; 50 ng/mL), ionomycin (1 μg/mL), and Brefeldin A (BFA; 1x) were added to stimulated PBMCs and BFA alone was added to unstimulated PBMCs for the final 6 hours of culture. All PBMC suspensions were then washed with Wash Buffer (Dulbecco’s Phosphate Buffered Saline (DPBS) with 1% FBS) and then incubated with fixable viability stain in the dark for 10 minutes at RT to label non-viable cells. PBMC were next suspended in 1:1 TruStain FcX blocker (Biolegend) and purified mouse IgG (BioRad) for 10 minutes at RT and then washed. PBMC were stained for surface antigens in Wash Buffer and Brilliant Stain Buffer Plus in the dark for 30 minutes at RT and then washed with Wash Buffer. PBMC were then fixed with 1.6% paraformaldehyde in DPBS for 20 minutes, washed, and permeabilized with FoxP3 Transcription Factor Staining Buffer (eBioscience) for 20 minutes at RT. PBMC were stained for intracellular antigens in the Transcription Factor Staining Buffer for 40 minutes at RT. Before acquisition, samples were washed three times and resuspended at 2.5 × 10^6^ cells/mL in Wash Buffer. Data was collected on a Cytek Aurora cytometer (5 Laser; SpectraFlo V3.1.2) and raw spectral data was ‘unmixed’ as previously described (*66*). PBMC stimulation and staining reagents, including fluorescent-tagged antibodies, and concentrations are outlined in **table S30**.

### Data Analysis

For each assay, measurements were taken from distinct samples. An exception to this was Olink proteomics, whereby three proteins had repeated measurements. This is detailed under the Plasma Proteomics section.

### Plasma proteomics

Protein abundance values were reported as log2 normalized protein expression (NPX) by Olink. Data were reviewed for overall quality prior to analysis. For cross-batch data normalization, bridge offsets were determined for each batch and each analyte separately by taking the median of the per-sample NPX differences between the later batch result and the earliest (reference) batch result for the 12 bridging controls. Offsets were subtracted from the analyte measurements of all samples in the later batch to obtain the normalized NPX values. Repeated assays (TNF, IL-6, and CXCL8) with non-significant Spearman correlations (P > 0.05) across panels and batches were excluded from further analysis. Proteins whose NPX values > 2 standard deviations from mean when comparing healthy samples from the first and last batches were excluded from analysis.

Ordinary least squares linear regression models were used to detect differences between groups, utilizing the *stats* package in R. Models were adjusted for age, sex, BMI (ARI vs. CON1) or age covariates (ERA vs. CON1). The Storey-Tibshirani procedure was employed for multiple testing correction, with q-values reported. Repeated assays (TNF, IL-6, and CXCL8) were treated as distinct measurements and therefore analyzed separately, as recommended by Olink. For each comparison, proteins were excluded from the analyses if > 50% of NPX values were below the limit of detection in either group. Longitudinal modeling of abundance changes associated with time to clinical RA are described in the Longitudinal Modeling section below. Geneset over- representation analysis was performed on significant proteins for a custom collection of genesets (MSigDB Hallmark v7.2, KEGG v7.2 and Reactome v7.2) using WebGestaltR (v0.4.6, method = ORA, 10^4^ permutations) with all protein coding genes as the background geneset. P values were calculated using hypergeometric tests, and adjusted using the Benjamini-Hochberg method.

### Flow Cytometry

#### Data Preprocessing, Clustering and Differential Abundance

Data from each panel were processed and analyzed independently. After spectral unmixing and manual compensation adjustments, cytometry data underwent pre-processing to remove technical artifacts, exclude doublets, and eliminate dead cells using a quadratic discriminant analysis model trained on manually gated data. A logicle transformation was applied to all fluorescent channels. Pre-trained machine learning models developed from healthy samples based on the Cyanno framework (*69*) were applied to facilitate cell type identification. The major populations (T, B, Myeloid, NK cells) were then subsetted from the corresponding target panel and downsampled to 40,000 cells per sample for downstream unsupervised clustering analysis. Subsequent dimensional reduction, batch-level harmonization and clustering on clinical samples within each panel were performed using Scanpy (*70*). Leiden clustering was performed at following resolutions: B, NK, myeloid cells at 1.0 in (PS, PB, PT, PM panels) and T cells at 2.0 in (PT, PS panels). Cluster-level counts and frequencies within each major cell type were calculated per sample and panel. Cluster phenotypes were assessed by comparing marker median expression across clusters and UMAP visualization. Statistical analysis of cellular abundance is detailed in the Differential Abundance Analysis section.

### Intracellular Flow Cytometry

#### Data Preprocessing, Metaclustering, Positivity Cutoffs and Statistics

Unmixed spectral data was manually compensated and total single and live cells were gated in FlowJo v10.10 Software. The gated data consisting of all live singlet PBMCs within the experiment, was then downloaded and further processed with the R programming language (http://www.r-project.org) and Bioconductor (http://www.bioconductor.org) software. Data was transformed with an inverse hyperbolic sine (asinh) transformation with a cofactor of 220. Each marker was scaled to the 99.9th percentile of expression of all cells within an experiment. Total live B cells were isolated *in silico* based on lineage marker expression, and then over-clustered into 100 clusters using FlowSOM with all informative surface molecules as input. Clusters were then hierarchically clustered based on expression of B cell surface molecules and isotype and finally manually assigned to cell subsets, as described previously (*71*). Cytokine positive thresholds for B cells were defined by the 99.9th percentile of expression for each cytokine in unstimulated B cells within each experiment (*72*).

### scRNA

#### Preprocessing, cleaning, and label transfer

Data preprocessing for scRNA were conducted per a computational reproducibility framework so that the analysis steps can be reproduced in the Human Immune System Explorer (*73*). High quality cells were selected based on the following cutoffs: <10% mitochondrial reads, number of genes detected between 200 and 5,000, RNA unique molecular identifiers (UMIs)/cells between 1,500 and 750,000. Doublets were removed using Scrublet (*74*) with default settings. The transcript of the remaining cells were first normalized, and then followed by variable feature selection, scaling, PCA, UMAP embedding as described in Scanpy (*70*). Cell type label transfer was performed using the Celltypist model (*75*) generated from a recently released AIFI Immune Cell Atlas (*24*) (**fig. S10A-C**). Following the cell type prediction, a secondary manual step was taken to remove the remaining doublet cells. All 71 cell types were individually subsetted and corrected for batch- and subject-level variation using Harmony (*76*), followed by Leiden clustering. Subsequently, clusters with gene expression indicative of doublets and high doublet scores, resulted in exclusion of 6.7% of cells from further analysis. Clusters with distinct gene profiles within the predicted cell type were manually reviewed and assigned unique cell type labels to differentiate from predicted cell types.

#### Pseudobulk differential gene expression analysis

Differential gene expression (DEG) analysis between groups was conducted using DESeq2 (*77*). For each sample and cell type pair, gene transcripts were aggregated using the get_pseudobulk function from decoupleR (*78*). Sample-cell type pairs with fewer than 10 cells or 1,000 total gene counts, as well as genes expressed in fewer than 10% of cells or in less than one-third of samples, or with fewer than 15 counts across all samples, were excluded from downstream analysis. Rare cell types present in fewer than five samples per group were also omitted from DEG analysis. Pseudobulk counts for the remaining cell types were analyzed with DESeq2 to compare ARI and CON1 groups with the formula (counts ∼ age + sex + BMI + group). Additionally, batch effects, particularly from batch B182, which exhibited a relatively stronger effect compared to other batches across multiple cell types during initial quality control (**fig. S10D**), were adjusted for in the model. For DEG analysis of longitudinal changes associated with time to clinical RA, pseudobulk counts were transformed with variance stabilizing transformation (VST) before being analyzed using a linear mixed model with the lmerTest package (described below). The Storey-Tibshirani procedure was employed for multiple testing correction, with q-values less than 0.1 considered significant.

#### Differential abundance analysis

Differential abundance analysis of immune cell populations across cytometry and scRNA data was conducted by first extracting counts and frequencies for each cluster or cell subset from all samples. To avoid zeros in ratio calculations, a pseudocount of 1 was added to the raw counts. Adjusted frequencies were then transformed using the centered log ratio (CLR) to account for the compositional nature of cell frequencies, utilizing the *composition* package in R. A linear regression model was employed to compare CLR values between ARI and CON1, adjusting for age, sex, and BMI. Multiple hypothesis testing was controlled using the Benjamini-Hochberg method as the number of tests were small, with FDR < 0.1 considered significant. Longitudinal modeling of DA analysis is performed as described below.

#### Longitudinal modeling of progression to clinical RA

Longitudinal analysis of ARI progression to clinical RA across all modalities (proteomic, scRNA, flow cytometry) for DEG and DA analyses were conducted using a linear mixed model in R (lmer function in lmerTest package), with Satterthwaite p-value approximation. The model was adjusted for age and BMI at the time of clinical RA diagnosis with subjects as the random effect (intercept only). Given the distinct sex differences and the limited number of samples available at earlier time points (**fig. S1B**), the analysis was restricted to samples from 13 female subjects within 750 days of clinical RA diagnosis. The model formula was as follows:

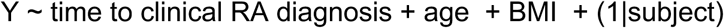

with the dependent variable (Y) being either the VST-transformed gene expression or the CLR-transformed cell frequency. The Storey-Tibshirani procedure was employed for multiple testing correction, with q-values less than 0.1 considered significant. For visualization, the effect sizes were annualized by multiplying by 365.

#### Longitudinal gene expression variability analysis

To investigate gene expression variability during ARI progression to clinical RA in comparison with the healthy aging process, we calculated intra-donor coefficients of variation (CVs) over time in ARI subjects who progressed to RA and matched controls (CON2) in the similar time frame, using the PALMO package (*79*). Pseudobulk gene counts, aggregated by Allen Institute for Immunology level 2 cell types, were normalized with variance stabilizing transformation. Intradonor CVs were computed using the *cvCalcBulkProfile* function, setting the time_column to "days to clinical RA" for converters and "days to the last visit" for CON2. CVs for each gene in each cell type were compared, and the percentage of genes with higher CVs in either group was reported.

#### Paired comparison between the last pre-symptomatic visit and clinical RA diagnosis visit

11 individuals whose samples are available at both pre-symptomatic visit and clinical RA diagnosis were included in the paired comparisons. (**Fig. 2E-I**) Pseudobulk and filtering steps were performed as described above. Paired DEG analysis was performed in DESeq2 (Counts ∼ subject + time point) to compare the gene expression between the two timepoints within the individual.

#### Gene program analysis

Supervised gene program analysis within and across cell types were conducted using Spectra (*26*). scRNA data of cross-sectional samples from ARI, ERA and CON1 were included in the analysis and downsample to the equal number of cells (6,366 cells) per sample to avoid sample bias in identifying the program. Global and cell-type-specific gene programs were obtained from Spectra as described and trimmed to match the major cell types in Fig S2C. RNA transcripts were normalized using Scran as suggested (*80*). Gene program modeling was performed using the est_spectra function in default setting. Single cell level factor scores were then extracted from the model results and scores above 0.001 were considered meaningful. The average scores per sample for each factor and cell type were then calculated and compared ARI and ERA to CON1 using a linear model, adjusted for age, sex, BMI, and batch effect. The Storey-Tibshirani procedure was employed for multiple testing correction, with q-values less than 0.05 considered significant.

#### T cell activation score

Activation scores for CD4 CM, core CD4 naive, and core CD8 naive cells were calculated by calculating the average expression from a manually curated gene set consisting of significantly up-regulated genes in the longitudinal model associated with T cell activation compared to the average expression of a randomly selected genes in the same size. (**Table S22**) The average scores for each sample were calculated and then the associations with time as ARI progressed to clinical RA were tested in linear mixed models (activation score ∼ days to clinical RA diagnosis + age + BMI + (1|subject)) as described above in the Longitudinal modeling section.

#### Pathway analysis

Gene Set Enrichment Analysis (GSEA) was performed as implemented in the FGSEA package to compare between the ARI and CON1 and with ARI progression to clinical RA based on the ranks genes list, where all detected genes were ordered by -log10(p)*sign(Log2 fold change). A custom collection of genesets was used, as described under Olink proteomics. The pathway enrichment *P* values were adjusted using the Benjamini-Hochberg method and pathways with adjusted *P* values < 0.05 were considered significantly enriched. Overlapping pathways are merged based on permutation test of independence by collapsePathways function in FGSEA package.

#### scRNA-seq B cell IgH Isotype Analysis

scRNA-seq data was used to estimate proportions of switched IgH isotope B cells in study samples. First, positive expression for each of the nine IgH genes (IGHM, IGHD, IGHG1-4, IGHA1-2, and IGHE) was defined per B cell as a normalized UMI count greater than or equal to 0.5 (normalized to 10,000 UMI per cell). Cells positive for either or both IGHM and IGHD were classified as ‘non-switched’ with isotype assignments of IGHM, IGHD, or IGHMD. The remaining cells were classified as ‘switched’ if positive for any other IGH gene. In cases of multi-positivity for switched genes, the positive gene located nearest the variable region in the genomic DNA was assigned as the isotype to be consistent with class-switch recombination biology (IgH Constant region ‘switched’ loci order after the variable region: IGHG3, IGHG1, IGHA1, IGHG2, IGHG4, IGHE, IGHA2). Both the switched status and isotype for cells without positive IgH gene expression were denoted as ‘undetermined’. After isotype classification, IgH germline transcription (GLT) level was defined as the normalized gene counts for those IgH loci downstream of the isotype-determining gene (**fig. S5**). As an example, IgG3^+^ isotype cells had quantifiable GLT from the IGHG1, IGHA1, IGHG2, IGHG4, IGHE, and IGHA2 genes. IgH constant region isotype gene GLT resulting in spliced and polyadenylated transcript is a well-established phenomenon in B cell isotype class-switching (*81–83*). Proportions of switched B cells were then compared between Memory B cell leiden clusters 8 and 9 using a Fisher’s Exact Test on the contingency table of B cell cluster versus switched status (R, fisher.test). scRNA-seq data was pseudobulked by B cell label and isotype to perform differential expression analysis of switched IgH genes (minimum 20 cells and 1,000 counts per sample). DESeq2 was used to compare ARI to CON1, with the model formula:

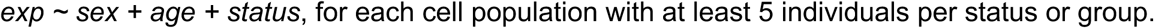

#### Nonnegative matrix factorization (NMF) analysis of the CD4 T cells

To identify changes in different T helper (Th) subsets in CD4 T cells, we performed NMF projection in total CD4 T cells based on a pre-computed weight matrix comprised of 12 NMF factors corresponding to T helper gene programs (*35*), which explained 24% of the total variance. Four NMF factors (NMF7-IFN, NMF8-CM, NMF9-Thymic-Emi, NMF10-Tissue) had low NMF scores across all cells in the dataset and NMF3-Naive is not relevant for memory cells, hence were omitted for the downstream analysis. We then subsetted the pan CD4 memory T cells and performed leiden clustering based on the remaining NMF factors at 0.4 resolution (**Fig. 5B**). Top differential genes of each cluster were identified using the Wilcoxon rank test in Scanpy. Relative frequency of clusters with the memory compartment for each sample was then calculated, CLR transformed, and modeled as described above in the Longitudinal Modeling section. C18 was a rare cluster (0.09% of CD4 memory cells) and present in only 68% of the samples, hence omitted from the longitudinal modeling.

#### Abatacept treatment response analysis

Gene expression matrices of bulk PBMC RNA-seq from 22 RA patients, both before and after abatacept (ABT) treatment, were obtained from a published study (*40*). DEG analysis was conducted using DESeq2 (*77*) to compare gene expression pre- and post-ABT treatment in responders and non-responders separately. Over-representation analysis was performed to test whether significant DEGs (FDR < 0.1) in each cell type from our longitudinal models of ARI progressing to clinical RA were enriched in ABT treatment responses of the responders and non-responders, respectively (Fig 5F). Cell type specific gene signatures with more than 10 genes were included in the analysis where hyperGeometric enrichment tests were performed in hypeR with background set to total expressed genes (*84*). The concordance of the effect sizes of the DEGs in RA progression and ABT responses were tested by McNemar’s Chi-squared Test. For all DEG analysis, the Storey-Tibshirani procedure was employed for multiple testing correction, with q-values less than 0.1 considered significant.

### TEA-seq

#### Data Preprocessing and Cell Labeling in TEA-Seq

Data preprocessing was performed as previously described (*68*). ADT and HTO count matrices from BarCounter were combined into a Mudata object (*85*). High quality cells were selected based on the following cutoff: > 250 genes per cell, between 500 and 20,000 RNA UMIs per cell, <10,000 ADT UMIs per cell, < 30% mitochondrial reads. One well (P1C2W6) was excluded from downstream analysis due to abnormally low RNA UMI counts and gene detection. ADT normalization is performed using the dsb package, which denoises ADT signals from background staining using empty droplets (*86*). RNA normalization, variable selection, scaling, and UMAP embedding were conducted using the standard Scanpy workflow (*70*). Fragment files for ATAC data were processed using the ArchR package (*87*). LSI reduced dimensions, UMAP and clusters generated from ArchR were then imported to the Mudata object for integrated analysis. 3-way weighted nearest neighbors and UMAP that incorporated ADT, RNA, and ATAC data were then generated using MUON (*85*). We performed unsupervised clustering within each major cell type (T, B, myeloid, and NK cells) and labeled the clusters based on the top differential expressed genes and ADTs (**fig. S7A**). In addition, label transfer based on RNA only was performed using the AIFI Immune Cell Atlas as described above in the scRNA section.

#### scATAC Peak Calling and Differential Peak Analysis

Peak calling for scATAC data in TEA-Seq was conducted using MOCHA (*88*). In brief, scATAC data were aggregated into cell type-sample pseudobulk matrices, and sample-specific 500 base pair tiles were identified. Consensus peaks were determined as regions that were open in more than 20% of samples. The zero-inflated Wilcoxon test implemented in getDifferentialAccessibleTiles function was then used to determine the differential accessible peaks between ARI and CON2 samples with settings of minimum median intensity (signalThreshold) = 14 and minimum difference in average dropout rates (minZeroDiff) = 0. FDR < 0.1 were considered significant. Peaks were annotated based on UCSC hg38 reference genome and regions within 2,000 base pairs upstream and 100 base pairs downstream of the transcription start site (TSS) were considered as promoter regions.

#### chromVAR Transcription Factor Activity Analysis

Single cell and pseudobulk transcription factor (TF) activities were inferred using chromVAR package (*89*). For single-cell TF activity, MOCHA-identified peaks were imported into the ArchR object, motifs were annotated using the CIS-BP database, and deviations were calculated using the addDeviationsMatrix function in ArchR. For pseudobulk-level analyses, TF deviations within CD4 naive T cells were calculated using the computeDeviations function, based on the MOCHA- identified pseudobulk sample peak matrix, and compared with GC-matched background peaks.

#### MOFA Analysis

Multi-Omics Factor Analysis (MOFA) was performed as described (*90*) to identify integrated factors across modalities in the TEA-Seq data. Normalized sample level pseudobulk ADT, RNA counts, TF deviation scores, and plasma proteins measured by Olink (described earlier) were used to train the MOFA model with the number of factors set to 6. Factor scores were extracted and compared between ARI and CON2 samples using a generalized linear model, controlling for age. Feature weights from each modality for factor 1 were also extracted to identify top-associated genes, TFs, ADTs, and plasma proteins.

#### Statistical Analysis

Specific statistical tests for each modality and analysis are described in the Data Analysis section above. All statistical tests were two-tailed.

## Acknowledgements

We thank the study participants for their valuable time and contributions to this research. We thank the Allen Institute founder, P.G. Allen, for his vision, encouragement, and support. We also thank all members of the Allen Institute for Immunology, in particular Maximilian Heeg and Anna Globig for critical review of the manuscript, Mansi Singh for critical review of analysis codes, the operations team for maintaining the productive research environment, and the Human Immune System Explorer (HISE) software development team for their support and dedication. This paper and the research behind it would not have been possible without HISE, a collaborative computational data analysis environment for life sciences research. Overview images created with BioRender.com.

## Funding

Funding was provided by the Allen Institute, NIH/NIAMS P30 AR079369 (KDD, VMH, MKD, KAK, LM and MLF), the University of Colorado Autoimmune Disease Prevention Center (KDD, MLF, CCS) and the William P. Arend Endowed Chair in Rheumatology Research (KDD). The content is solely the responsibility of the authors and does not necessarily represent the official views of the National Institutes of Health.

## Author Contributions

Conceptualization: ZH, MLF, MKD, KAK, HT, WW, FZ, DLB, TFB, VMH, AKS, GSF, KDD, TRT, MAG

Methodology: MCG, PR, EMD, ATH, RL, SRZ, DLB, PJS, TRT

Software: CML, LTG

Investigation: MCG, MLF, LL, PR, EMD, AGA, NAA, EAB, JHB, SB, CEC, MAC, MKD, CLF, JG, PCG, BCH, VH, ATH, EKK, KAK, KJL, CL, LM, RRM, BM, KN, AO, CP, CGP, VP, JR, CRR, CS, CCS, TJS, ES, HT, TT, VT, ZJT, JAS, MDS, MDW, FZ, DLB, VMH, KDD

Resources: CEG, ELK, LTG, QG, TP

Data curation: PV, MLF, BCH, LM, NT, UK, KDD

Formal analysis: ZH, MCG, PV, LYO, NTT, YH, SRZ, M-PP, UK, AKS, MAG Supervision: ZH, YH, AWG, TFB, XL, PJS, VHM, AKS, GSF, KDD, TRT, MAG

Project administration: CEB, LAB, AW Funding acquisition: AWG, TFB

Writing - original draft: ZH, MCG, AKS, TRT, MAG

Writing - review and editing: ZH, MCG, MLF, LYO, NTT, YH, PR, EMD, RL, FZ, TFB, XL, PJS, VMH, GSF, AKS, KDD, TRT, MAG

## Competing interests

KDD has received honorarium and reduced-cost biomarker assays from Werfen. KDD and MKD receive grant funding from Gilead Sciences. GSF, LYO, NTT, YH, CEB, XL and TRT receive grant funding from Eli Lilly. AWG serves on the scientific advisory boards of Arsenal Bio and Foundery Innovations and is a cofounder of TCura.

## Data and Materials Availability

Processed scRNA-seq data from human PBMCs generated will be deposited in GEO and released upon publication. Raw scRNA-seq fastq files will be deposited in dbGap and be publicly available on the date of publication. Abatacept treatment data from RA patients was derived from Iwasaki et al. (2024) and can be downloaded from Zenodo (https://zenodo.org/records/8250013).

## Code Availability

R and Python code used to perform analyses and generate figures is provided in GitHub (https://github.com/aifimmunology/ALTRA-manuscript/).

## Data Visualizations

The following tools are provided to facilitate data exploration and discovery:

Systemic inflammation in At-risk Individuals Advancing to Clinical Rheumatoid Arthritis https://apps.allenimmunology.org/aifi/insights/ra-progression/

1. ***Rheumatoid arthritis scRNA-seq UMAP Explorer***. Explore cell subsets and gene expression in a UMAP viewer to display an overview of the entire dataset. https://apps.allenimmunology.org/aifi/insights/ra-progression/vis/scrna-umap/
2. ***Rheumatoid arthritis TEA-seq Explorer***. Explore trimodal gene expression, surface protein expression, and chromatin accessibility from a subset of participants. https://apps.allenimmunology.org/aifi/insights/ra-progression/vis/tea-umap/
3. ***Rheumatoid arthritis Gene and Protein Expression Explorer***. Explore cross-sectional and longitudinal changes in gene expression and circulating protein abundances. https://apps.allenimmunology.org/aifi/insights/ra-progression/vis/gene-protein-explorer/

## Supplementary Materials

**Fig S1.**
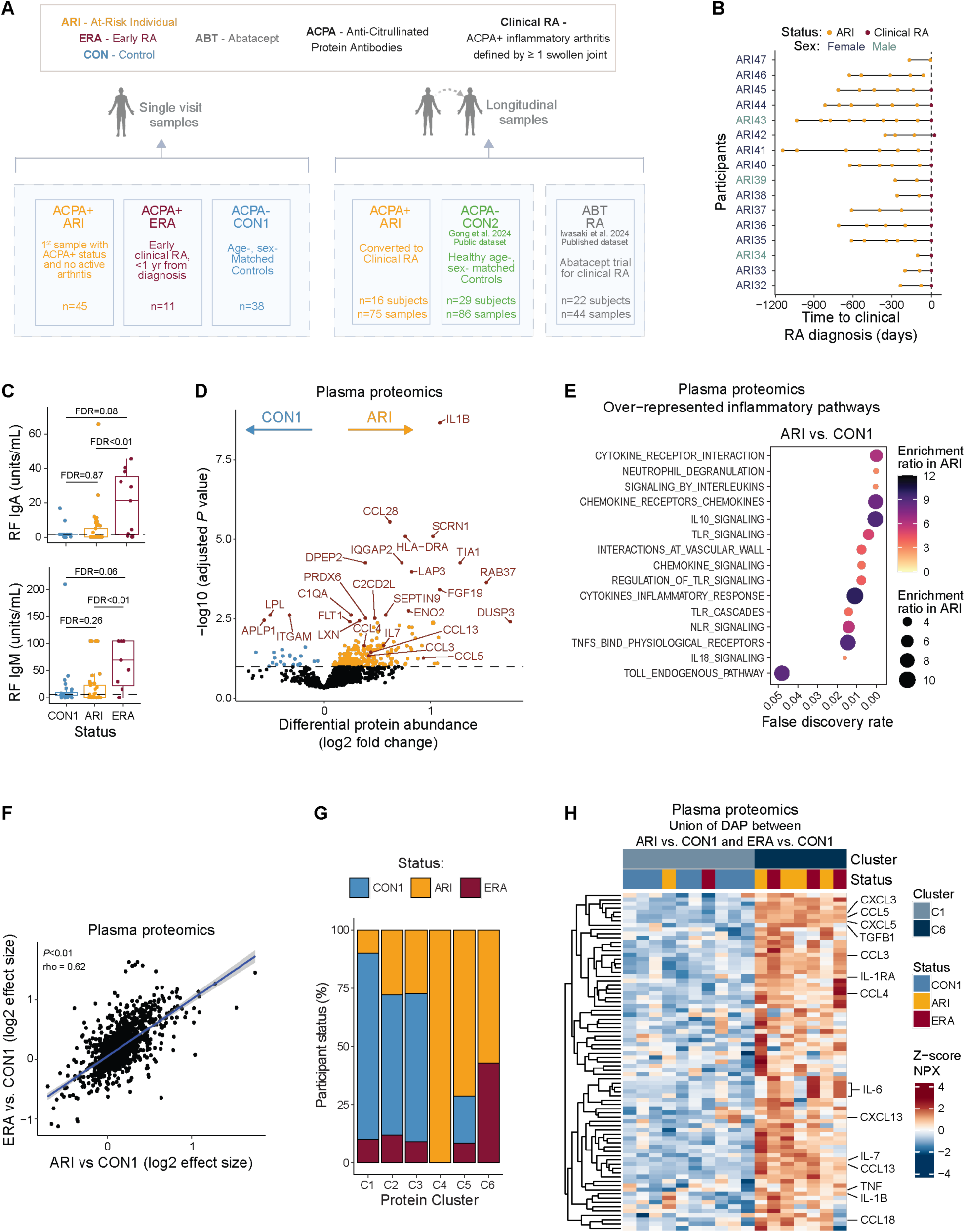
related to data figure 1. (A) Consort diagram of groups and data sets used in this study. (B) Longitudinal blood sampling frequency of ARI who progressed to clinical RA. Onset of clinical RA is denoted by dashed vertical line (time to clinical RA diagnosis = 0). Biological sex is denoted by participant ID color. (C) Baseline (initial) RF-IgA and RF-IgM levels from CON1, ARI, ERA. (D) Volcano plot of baseline differential plasma protein abundance elevated in ARI (orange) or CON1 (blue). Each dot represents a single protein assayed. The 20 proteins with the smallest *P* values, in addition to select inflammatory proteins, are noted in red. (E) Over-represented inflammatory pathways (MSigDB Hallmark, KEGG, Reactome) in ARI. Enrichment ratios are shown by color and size. (F) Comparison of differential protein abundance between ARI vs. CON1 and ERA vs. CON1 (Spearman ρ = 0.62). (G) Number of ARI, ERA, CON1 participants comprising each protein cluster from Fig. 1C. (H) Z-scored NPX of differentially abundant inflammatory mediators between clusters Prot-C1 and Prot-C6. Columns indicate participants. Select proteins (rows) are labeled. Boxplots show median (centerline), first and third quartiles (lower and upper bound of the box) and whiskers show the 1.5x interquartile range of data. *P* values were calculated by Kruskal-Wallis followed by Dunn’s post hoc testing (C), linear regression models (D), or hypergeometric tests (E). Nominal *P* value is indicated for (F). FDR values are indicated for remaining panels.

**Fig. S2.**
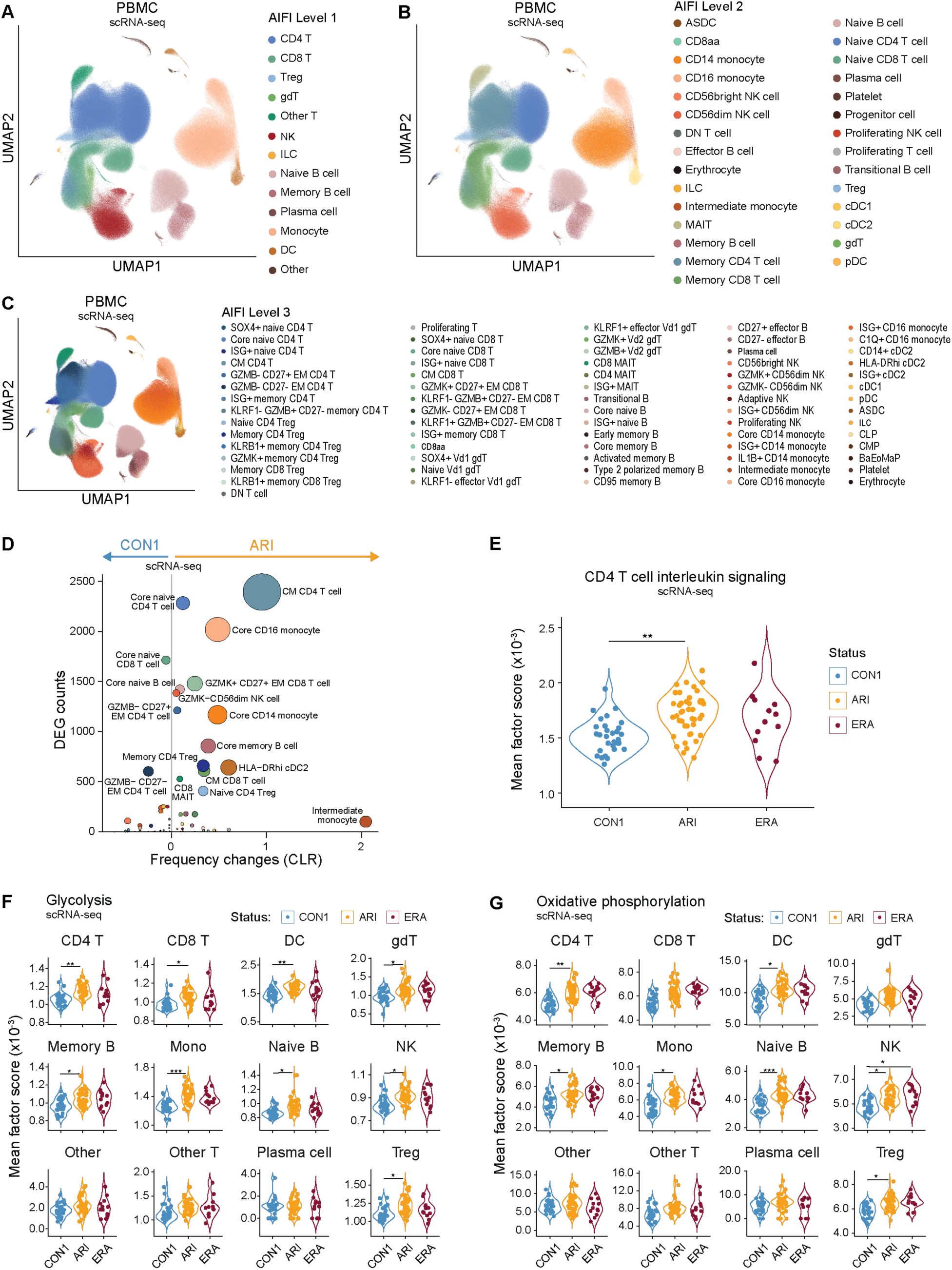
related to data figure 1. (A-C) UMAPs of cell subsets labeled using the Allen Institute for Immunology Immune Cell Atlas at levels 1 (A), 2 (B), and 3 (C). (D) Comparison DEG number with the change in frequency (centered-log ratio (CLR) transformation) between ARI and CON1 at level 3 cell types. Bubble size corresponds to the aggregate score calculated by [-log(padj CLR frequency changes) x total number of DEGs]. (E-G) Spectra factor scores for interleukin signaling in CD4 T cells (E), and oxidative phosphorylation (F) and glycolysis (G) from ARI, ERA, CON1 across all level 1 cell types. *P* values were calculated using linear regression modeling (D-G). For (E-G), all pairwise comparisons were tested and FDR values are indicated for those that are significant. *FDR < 0.05; **FDR < 0.01.

**Fig. S3.**
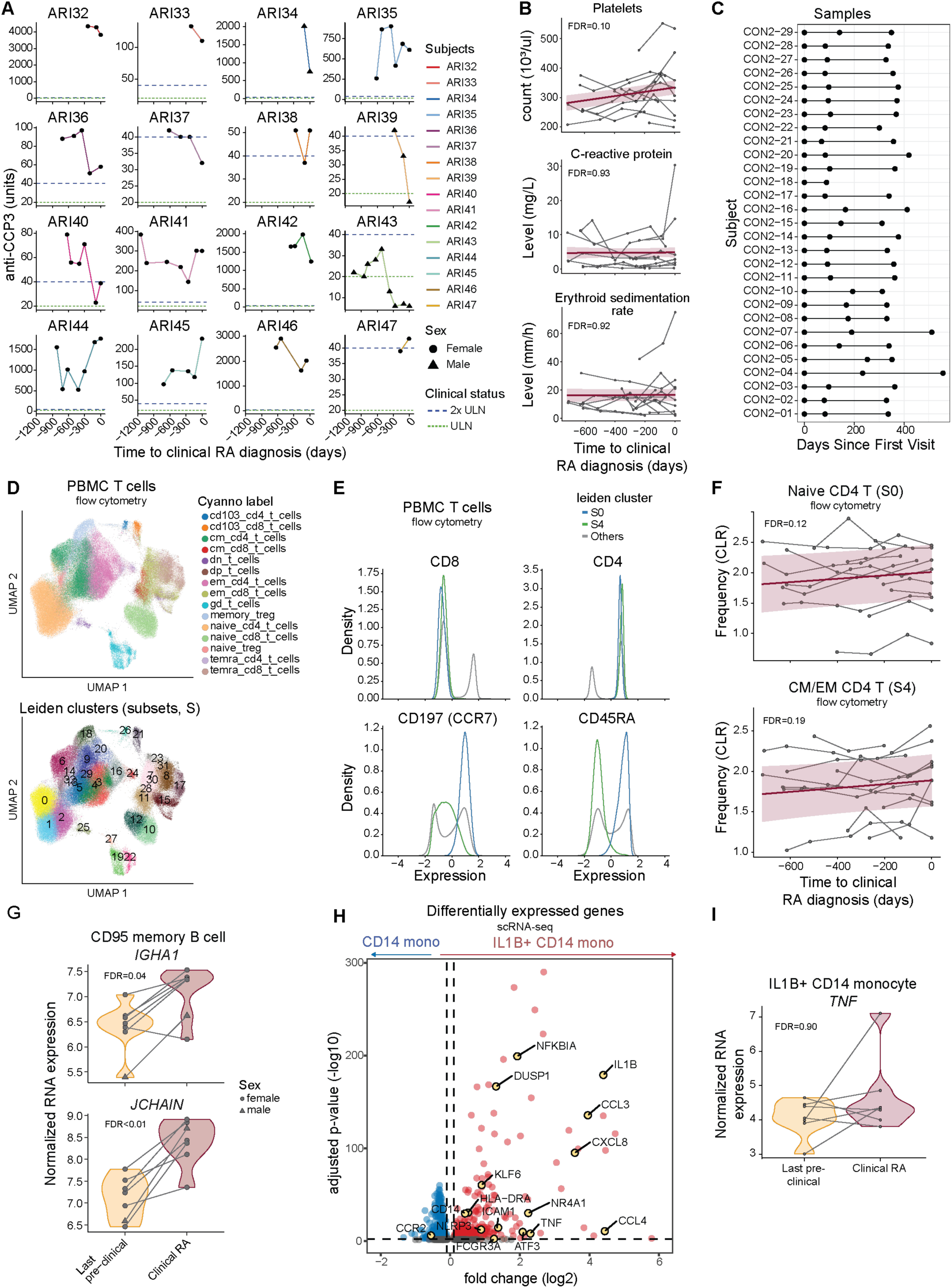
related to data figure 2. (A) Longitudinal anti-CCP3 serum levels in ARI who progressed to clinical RA. Each plot represents a different participant, and biological sex is indicated by point shape. Dashed horizontal lines indicate the upper limit of normal (ULN; 20 units; green) and 2x ULN (40 units; blue). (B) Clinical lab features in ARI who progressed to clinical RA. Each participant’s longitudinal series is connected by a gray line, with a group trendline and 95% confidence interval in purple. (C) Longitudinal sample inclusion from CON2 for comparison of intra-donor coefficients of variation in Fig. 2B. (D-F) PBMC were analyzed by flow cytometry and clustered into subsets (S) for abundance comparisons. Significant subsets were manually reviewed and annotated. (D) UMAPs annotated for Cyanno cell type labels (top) and clustered subsets (bottom). (E) T cells markers in S0 and S4 compared to all other subsets. (F) Centered log-ratio (CLR)-transformed frequency changes of subsets annotated as naive CD4 and central memory/effector memory (CM/EM) CD4 T cells over time as ARI progress to clinical RA. (G) RNA expression of IGHA1 and JCHAIN in CD95 memory B cells. Paired donor samples from their last pre-symptomatic and diagnosis of clinical RA visits are connected by lines. (H) Volcano plot comparing genes with elevated expression in IL1B+ CD14 monocytes (red) vs. core CD14 monocytes (blue). (I) Normalized RNA expression of TNF in IL1B+ CD14 monocytes, as in (G). *P* values were calculated using linear mixed models (B-C), paired Wald test (G,I), or Wilcoxon rank-sum test (H). FDR values are indicated for all panels.

**Fig. S4.**
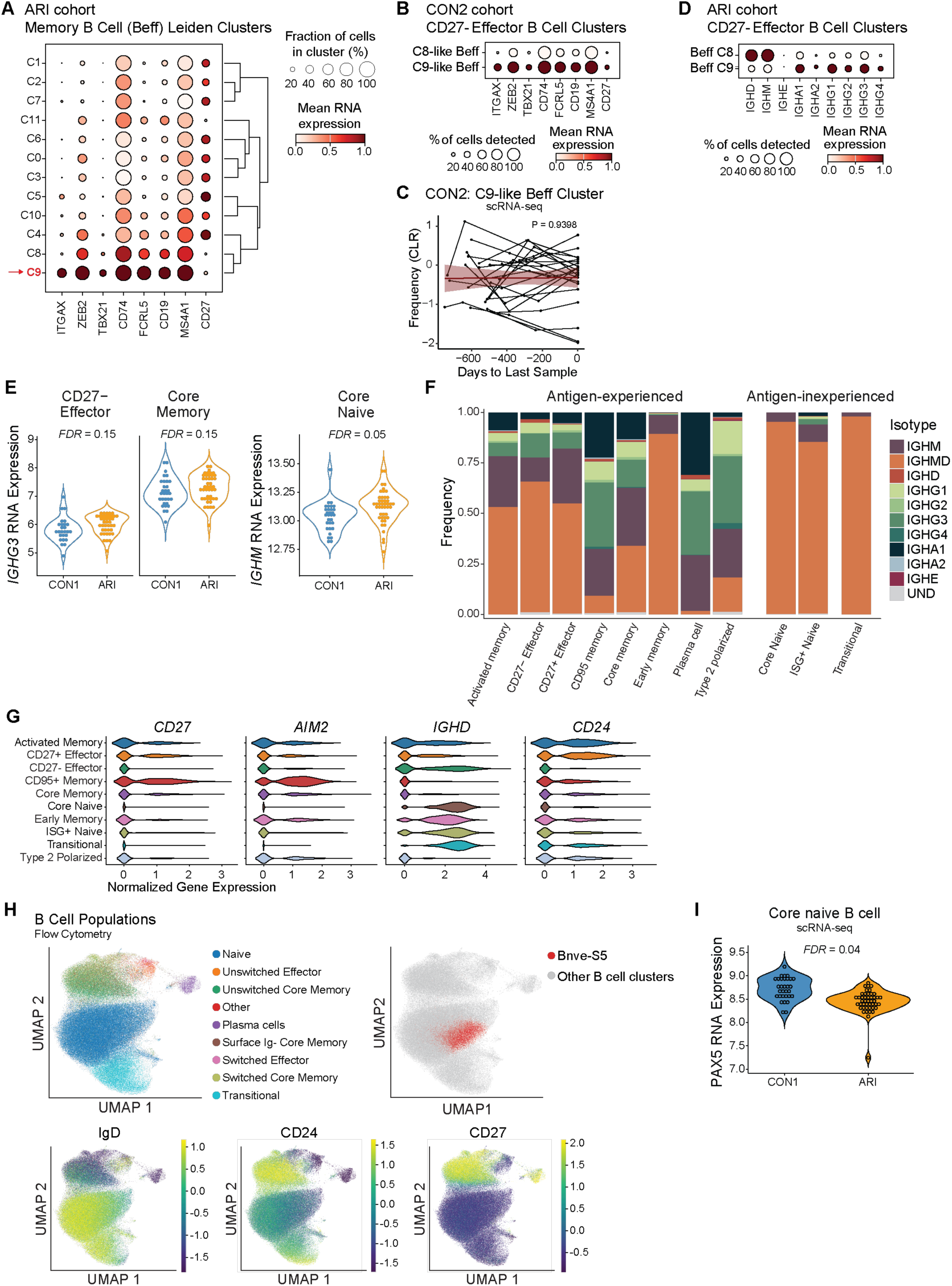
related to data figure 3. (A) Subset-defining expression among ARI memory B cell leiden clusters from scRNA-seq. (B) Subset-defining gene expression for CD27-effector B cell identity clusters. (C) Longitudinal centered log-ratio (CLR) transformed frequencies for Beff C9-like cluster in CON2 over a 2-year span. Each participant’s longitudinal series is connected by a gray line, with a group trendline and 95% confidence interval in purple. (D) IGH gene expression levels for Beff-C8 and Beff-C9 from ARI. (E) *IGHG3* RNA expression by core memory (P=0.02; FDR=0.15) and CD27-effector (P=0.004; FDR=0.15) B cells and *IGHM* expression by naive B cells (P=0.02; FDR=0.05) of ARI and CON1. (F) B cell IgH isotype composition, as frequency within population, for all subsets. (G) CD27, AIM2, CD24 and IGHD gene expression by B cell subsets in ARI. (H) Flow cytometry UMAP plots for B cells showing population labels determined by Cyanno model-based approach (top left), Bnve-S5 cells (top right, red dots), and overlaid subset-defining marker expression (bottom). (I) PAX5 gene expression in naive B cells from ARI and CON1. *P* values were calculated using linear mixed models (C), Wald test in DESeq2 with Storey-Tibshirani procedure (E), and Wilcoxon rank-sum test (I). Nominal *P* value is indicated for (C). FDR values are indicated (E,I).

**Fig. S5.**
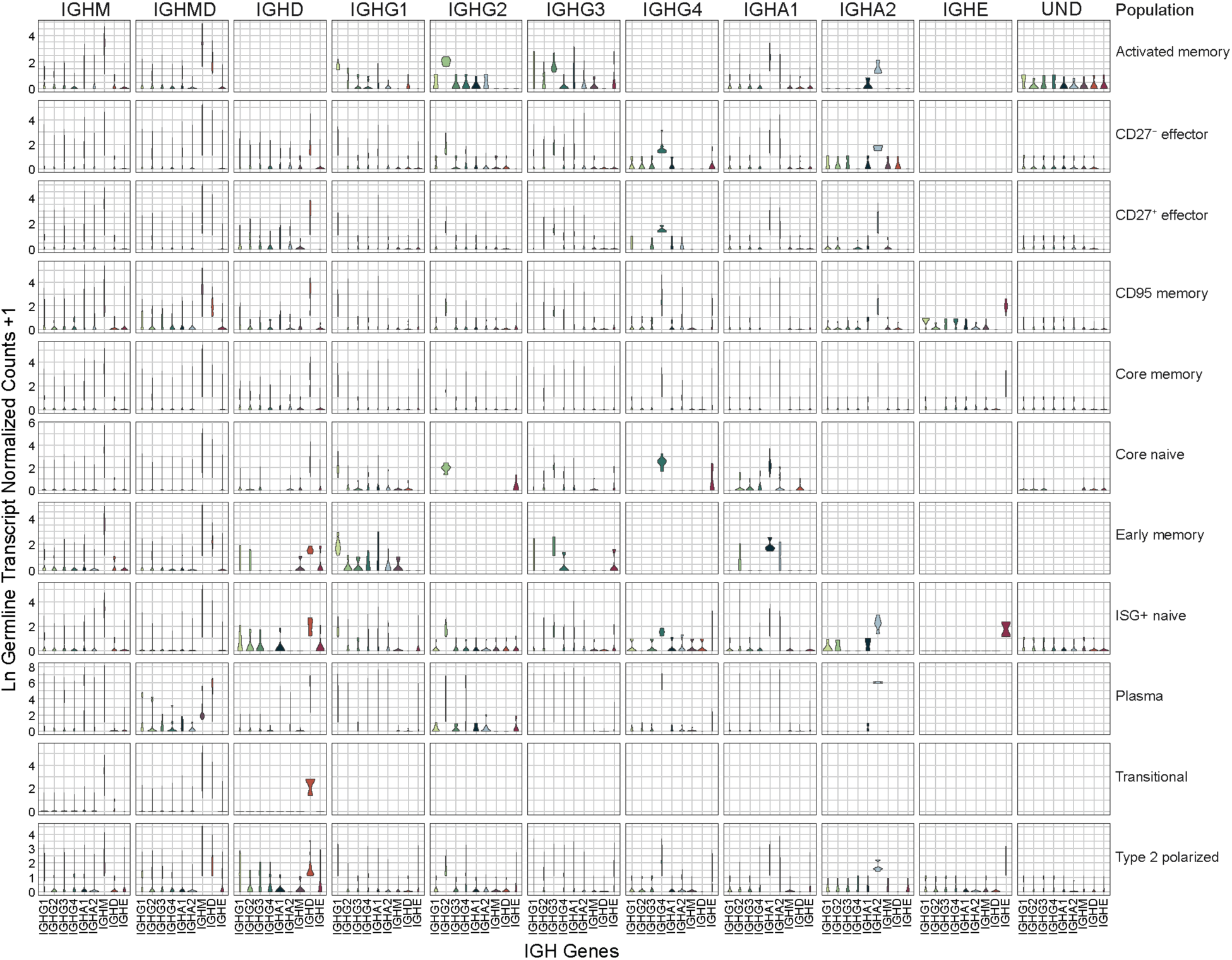
related to data figure 3 and methods. Log1p-transformed IGH gene germline transcription (GLT) normalized counts for each B cell isotype and subset in scRNA-seq data.

**Fig. S6.**
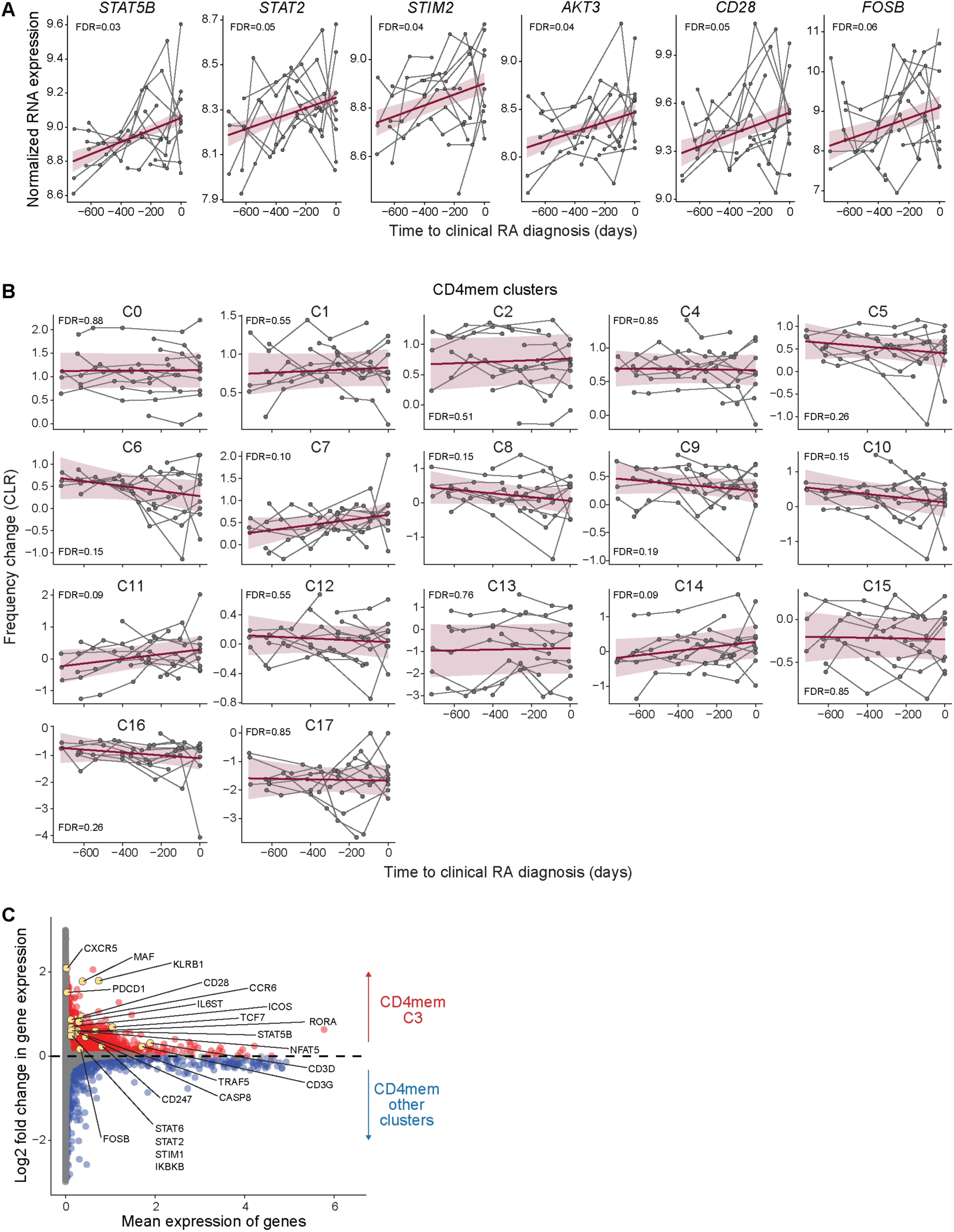
related to data figure 4. (A) RNA expression of select genes associated with activation in central memory (CM) CD4 T cells as ARI progress to clinical RA. Genes were selected based on Fig. 4A. Each participant’s longitudinal series is connected by a gray line, with a group trendline and 95% confidence interval in purple. (B-C) CD4mem Leiden clusters were derived from non-negative matrix factorization (NMF)-projected CD4 reference gene weights from Yasumizu *et al*. onto CD4mem T cells (see Fig. 4C). (B) Centered log-ratio (CLR)-transformed frequency for each cluster is shown over time as ARI progress to clinical RA. Group trendlines were determined as in (A). (C) Comparison of RNA expression in cluster CD4mem-C3 (red) vs all remaining clusters (blue) over time as ARI progress to clinical RA, with mean expression of each gene. *P* values were calculated using linear mixed models (A-B). FDR values are indicated for all panels.

**Fig S7.**
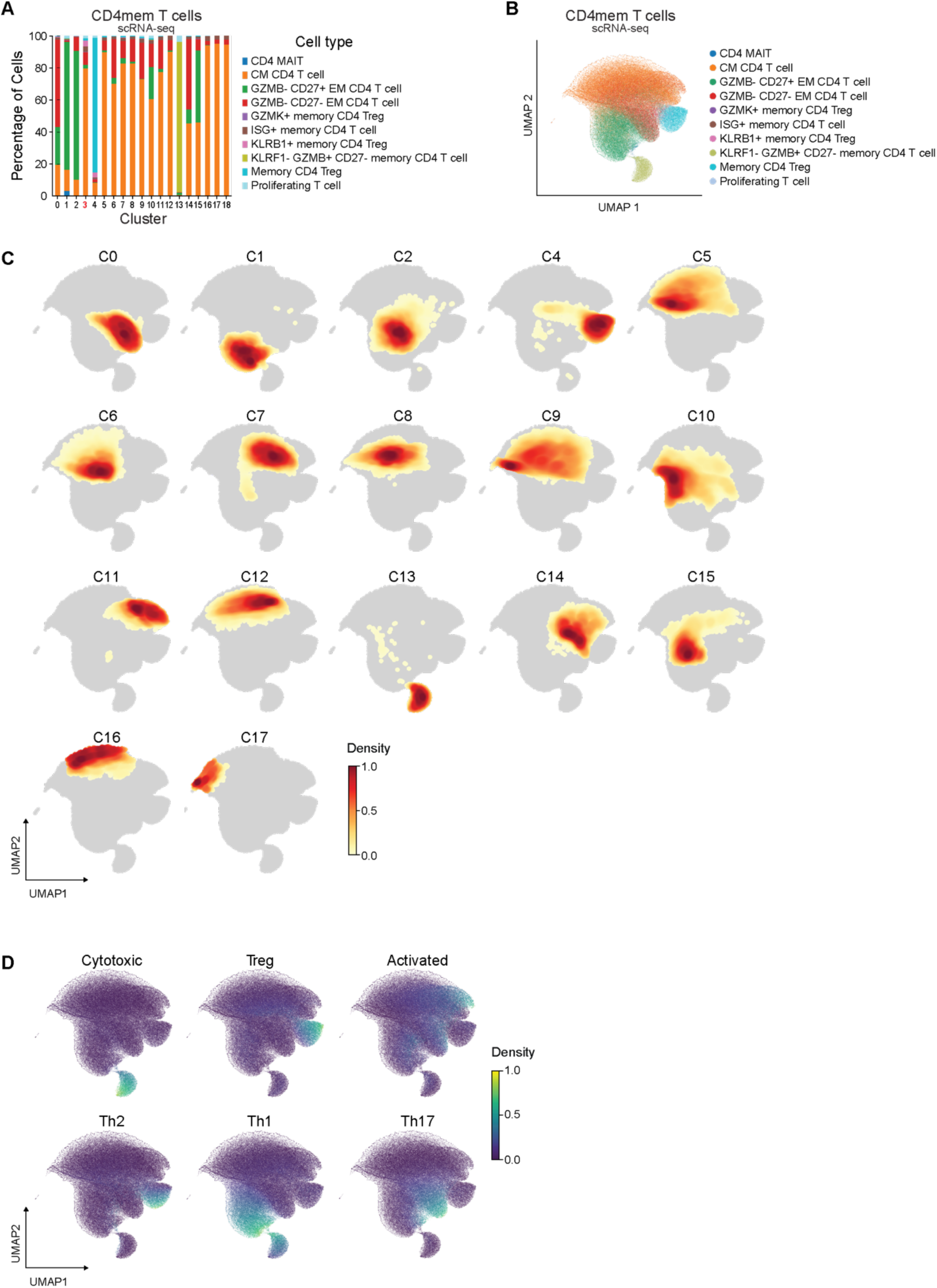
related to data figure 4. (A-D) CD4mem T clusters were derived as in fig. S6B. Quantitation (A) and UMAP (B) by Allen Institute for Immunology Immune Cell atlas level 3 labels. (C) UMAP density plots for each cluster. CD4mem-C3 is shown in Fig. 4H. (D) CD4mem T cells expressing polarized gene programs are distinguished based on the NMF projection using a pre-computed weight matrix of CD4 T cell population from Yasumizu *et al*. The Tfh density is shown in Fig. 4H.

**Fig. S8.**
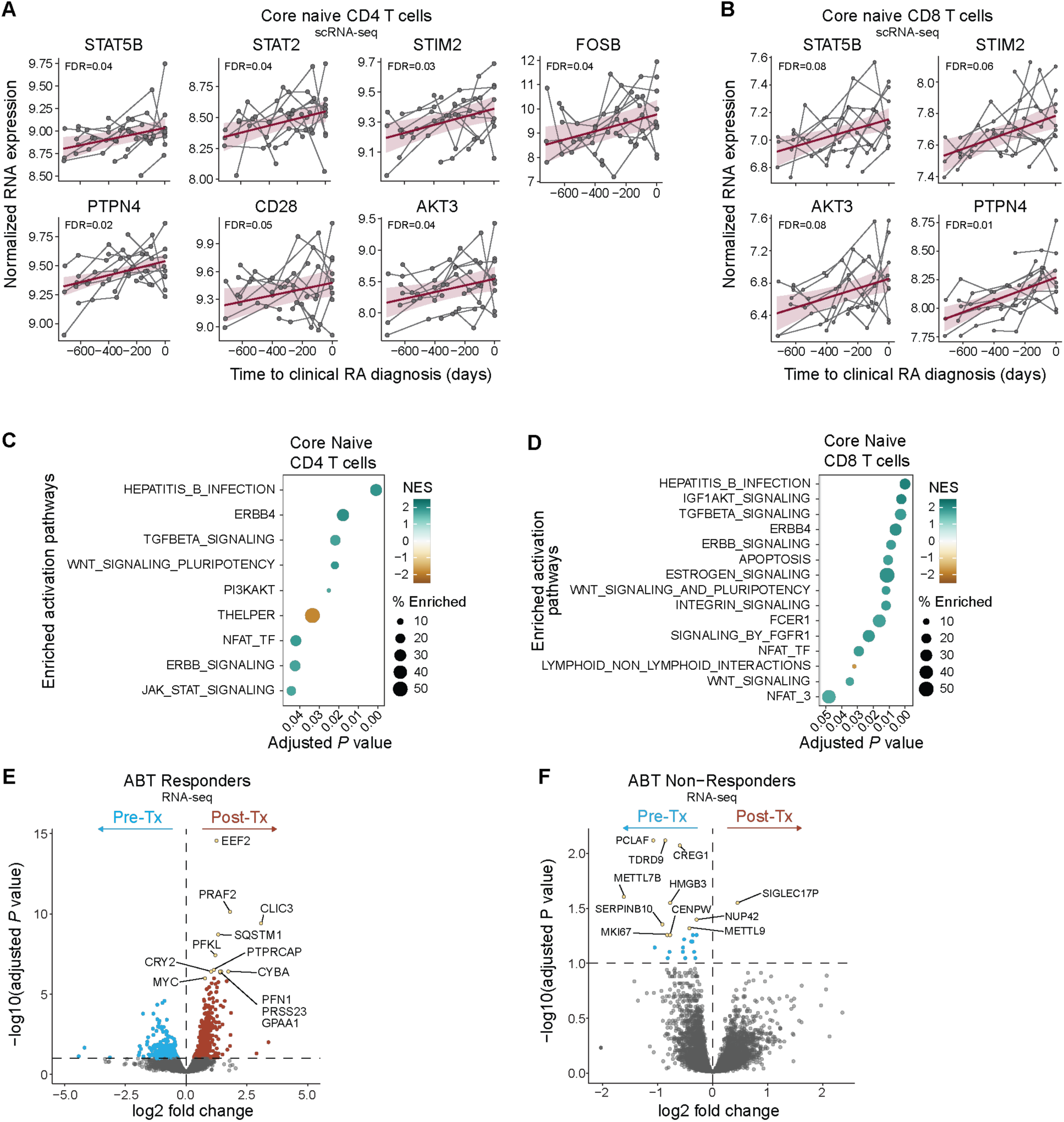
related to data figure 5. (A,B) Changes in RNA expression of select genes associated with T cell activation over time as ARI progress to clinical RA in core naive CD4 (A) and CD8 (B) T cells. Genes were selected based on Fig. 5A and Fig. 5C. Group trendlines, derived from linear mixed models, are shown with a purple line with 95% confidence interval in the shaded area. (C,D) Enriched pathways associated with T cell activation in core naive CD4 (C) and CD8 (D) T cells over time as ARI progress to clinical RA. Normalized enrichment scores (NES), by GSEA, are shown. (E,F) Volcano plots, derived from reanalysis of Iwasaki et al. (Fig. 5E), showing the differential expressed genes in responders (D) and non-responders (E) before and after abatacept (ABT) treatment. *P* values were calculated using linear mixed models (A-B) or Wald test (E-F). FDR values are indicated for all panels.

**Fig. S9.**
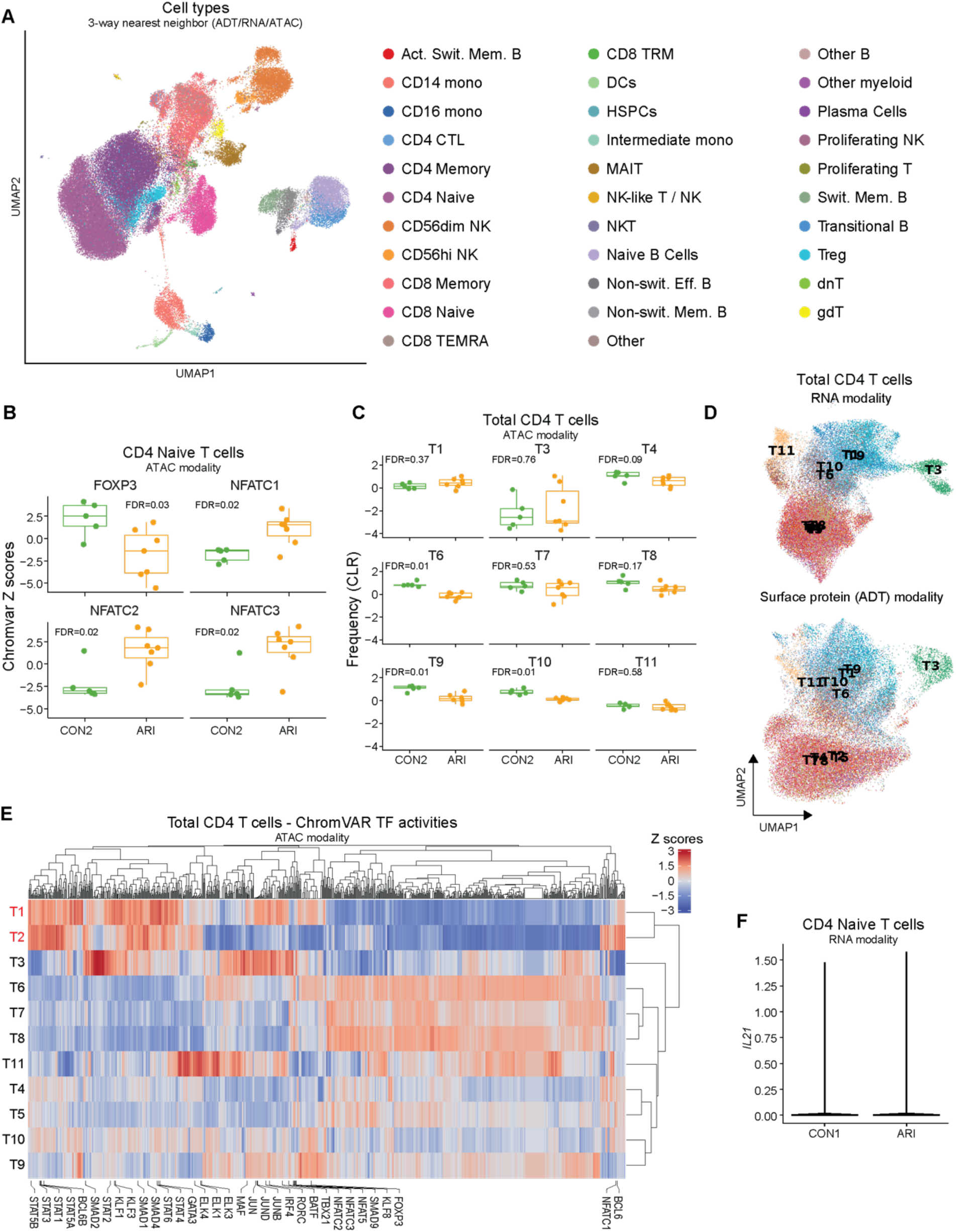
related to data figure 6. (A) Three-way weighted nearest neighbor UMAP of TEA-seq data incorporating surface protein, transcript, and chromatin accessibility, colored by cell type label. (B) ChromVAR Z scores of selected transcription factor activities in naive CD4 T cells. (C) CLR frequency of CD4 T cells ATAC clusters. (D) UMAP of CD4 T cells in RNA modality (top) and surface protein (ADT) modality (bottom). (E) Heatmap of ChromVAR TF activity scores of 870 TFs among clusters, scaled by column. Selected TFs related to T cell activation and differentiation are labeled. (F) Normalized RNA expression of IL21 in ARI and CON1. Boxplots show median (centerline), first and third quartiles (lower and upper bound of the box) and whiskers show the 1.5x interquartile range of data. *P* values were calculated using the Wilcoxon rank-sum test (B-C). FDR values are indicated (B-C).

**Fig. S10:**
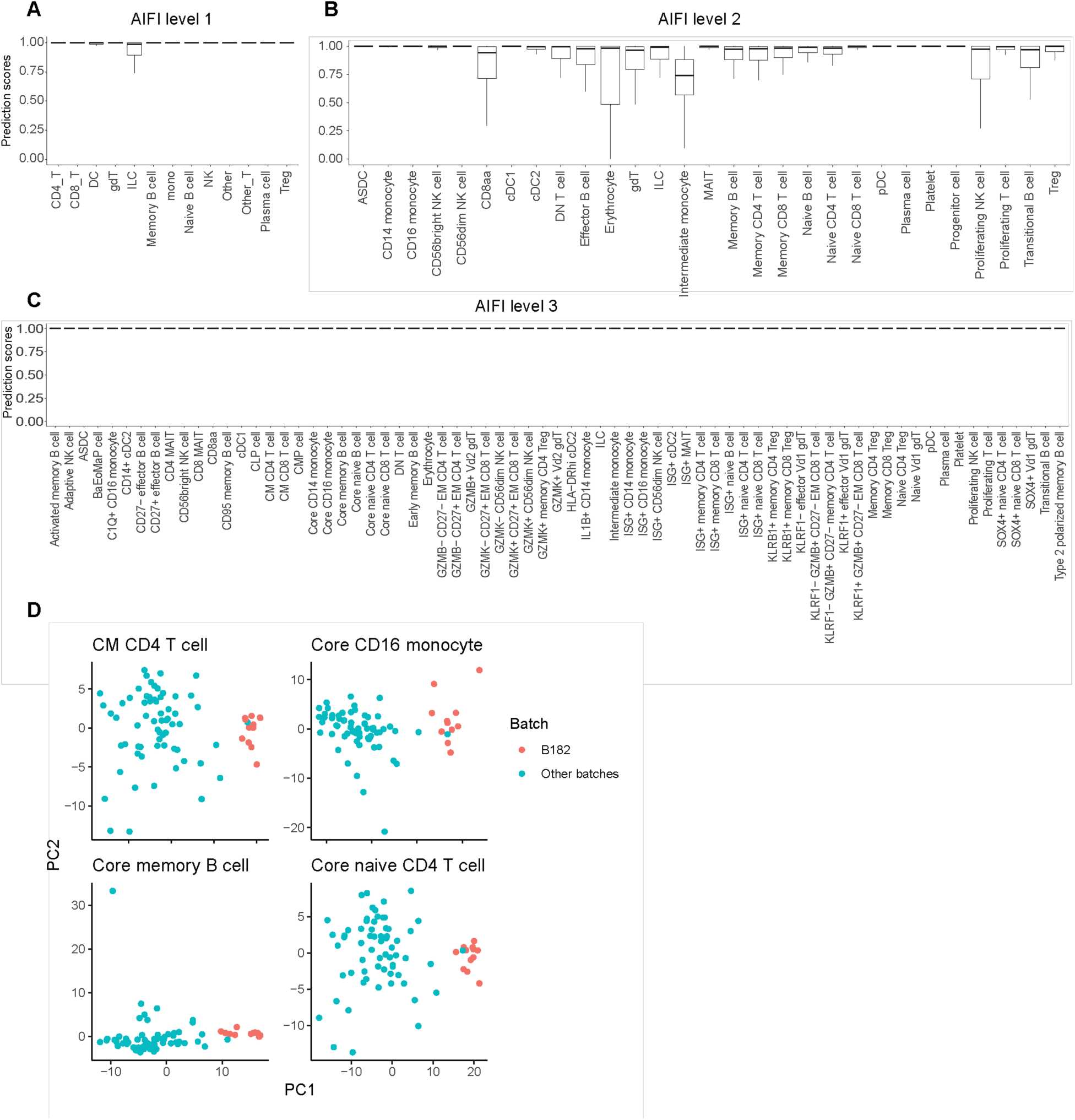
related to methods. Prediction scores of the CellTypist model generated from Allen Institute for Immunology (AIFI) Immune Cell Atlas cell types for level 1 (A), level 2 (B), and level 3 (C). Boxplots show median (centerline), first and third quartiles (lower and upper bound of the box) and whiskers show the 1.5x interquartile range of data. (D) Principal component analysis of pseudobulk scRNA data for CM CD4 T cells, core memory B cells, and core CD16 monocytes. Each dot represents a sample, colored by whether the sample was processed in batch 182 or other batches in the dataset.

**Supplementary Table S2:** Summary of baseline cohort information.

| Variable | ACPA-<br>CON1<br>(n = 38) | ACPA+<br>ARI<br>(n = 45) | ACPA+<br>ERA<br>(n = 11) | P value<sup>1</sup> |
| --- | --- | --- | --- | --- |
| Age at sample collection, mean years (SD) | 56 (16) | 57 (16) | 49 (9) | 0.23 |
| Sex: Female, n (%) | 34 (89%) | 36 (80%) | 10 (91%) | 0.52 |
| Sex: Male, n (%) | 4 (11%) | 9 (20%) | 1 (9%) | - |
| Ethnicity: Non-Hispanic origin, n (%) | 29 (76%) | 43 (96%) | 8 (73 %) | 0.02 |
| Race: White, n (%) | 32 (84%) | 37 (82%) | 7 (64%) | 0.27 |
| BMI, mean kg/m<sup>2</sup> (SD) | 28 (6) | 27 (5) | 30 (8) | 0.66 |
| Met ACR/EULAR 2010 criteria, n (%) | - | - | 11<br>(100%) | - |
| C-reactive protein, median mg/L (IQR) | 1.6 (0.6-3.4) | 1.5 (0.9-4.7) | 3.3 (1.0-10) | 0.39 |
| Erythrocyte sedimentation rate, median<br>mm/h (IQR) | 9 (4-13) | 11 (7-20) | 20 (11-33) | 0.07 |
| Shared epitope present, n (%)<sup>2</sup> | 13 (38%) | 18 (40%) | 6 (55%) | 0.69 |
| Serum ACPA, median (IQR) | 6 (6-6) | 62 (44-195) | 560 (230-1101) | <0.01<br>(<0.01 Con vs. ARI;<br><0.01 Con vs. ERA) |
| Serum RF IgM, median (IQR) | 5 (5-9.7) | 0.9 (0.3-23) | 69 (23-105) | 0.01<br>(<0.01 ARI vs. ERA;<br>0.06 Con vs. ERA) |
| Serum RF IgA, median (IQR) | 1.7 (1.7-1.7) | 0.3 (0.3-5.2) | 21 (1.4-35) | <0.01<br>(<0.01 ARI vs. ERA;<br>0.08 Con vs. ERA) |

<sup>1</sup>P values were calculated using the Kruskal-Wallis test for continuous variables. For $P < 0.05$ , a Dunn's post-hoc test with Bonferroni correction was performed and significant results are indicated in parentheses. P values were calculated using the Chi-squared test for discrete variables.
<sup>2</sup>Shared epitope present at any *HLA-DRB1* \*01:01, \*01:02, \*04:01, \*04:04, \*04:05, \*04:08, \*04:09, \*04:10, \*04:13, \*10:00 alleles. We were unable to measure 4 CON1 participants, and these were excluded from the percent calculation.

**Supplementary Table S3:** Longitudinal summary of ACPA+ ARI who progressed to clinical RA.

| Variable | All participants (n = 16) |  |  | Female only (n = 13) |  |  |
| --- | --- | --- | --- | --- | --- | --- |
|  | Baseline Visit | IA Onset | P value<sup>1</sup> | Baseline Visit | IA Onset | P value<sup>1</sup> |
| Age at sample collection, mean years (SD) | 51 (16) | 52 (16) | <0.01 | 46 (14) | 47 (14) | <0.01 |
| Sex: Female, n (%) | 13 (81%) | - | - | 13 (100%) | - | - |
| Ethnicity: Non-Hispanic origin, n (%) | 15 (94%) | - | - | 12 (92%) | - | - |
| Race: White, n (%) | 11 (69%) | - | - | 9 (69%) | - | - |
| BMI, mean kg/m<sup>2</sup> (SD) | 28 (5) | 28 (5) | 0.49 | 27 (6) | 28 (6) | 0.07 |
| Met ACR/EULAR 2010 criteria, n (%) | - | 12 (75%) | - | - | 10 (77%) | - |
| C-reactive protein, median mg/L (IQR) | 2.8 (0.8-6.2) | 2.3 (1.2-3.3) | 0.98 | 3.3 (0.8-8.4) | 2.9 (1.2-4.1) | 1.00 |
| Erythrocyte sedimentation rate, median mm/h (IQR) | 12 (9-25) | 14 (8-23) | 0.71 | 12 (9-23) | 13 (9-21) | 0.67 |
| Shared epitope present, n (%)<sup>2</sup> | 6 (38%) | - | - | 5 (38%) | - | - |
| Serum ACPA, median (IQR) | 115 (49-1581) | 171 (42-875) | 0.11 | 133 (79-1557) | 231(51-1242) | 0.29 |
| Serum RF IgM, median (IQR) | 15 (0.3-34) | 15 (3.2-35) | 0.22 | 16 (0.3-30) | 16 (3.8-50) | 0.11 |
| Serum RF IgA, median (IQR) | 0.3 (0.3-5.6) | 1.7 (0.3-2.7) | 0.92 | 0.3 (0.3-5.2) | 1.7 (0.3-4.1) | 0.94 |
| Average duration in the study, days (min, max) | - | 467.9 (106-717) | - | - | 497.8 (163-717) | - |

<sup>1</sup>P values were calculated using the paired Wilcoxon rank-sum test for continuous variables. P values were calculated using the Fisher exact test for discrete variables.
<sup>2</sup>Shared epitope present at any *HLA-DRB1* \*01:01, \*01:02, \*04:01, \*04:04, \*04:05, \*04:08, \*04:09, \*04:10, \*04:13, \*10:00 alleles.

**Supplementary Table S13:** Summary of healthy controls (CON2) with longitudinal sampling.

| Variable | CON2 baseline visit (n=29) | CON2 last visit (n=29) | *P* value<sup>1</sup> |
| --- | --- | --- | --- |
| Age at sample collection, mean years (SD) | 47.2 (14.2) | 48.0 (14.2) | <0.001 |
| Sex: Female, n (%) | 22 (76%) | - | - |
| Ethnicity: Non-Hispanic origin, n (%) | 28 (97%) | - | - |
| Race: White, n (%) | 23 (79%) | - | - |
| BMI, mean kg/m<sup>2</sup> (SD) | 27.4 (5.0) | 28.2 (4.4) | 0.76 |
| Met ACR/EULAR 2010 criteria, n (%) | 0 | 0 | - |
| C-reactive protein, median mg/L (IQR) | 1.9 (1.1-3.0) | 1.5 (0.7-2.0) | 0.18 |
| Erythrocyte sedimentation rate, median mm/h (IQR) | 2 (2-9) | 2 (2-9) | 0.25 |
| Serum ACPA, median (IQR) | 0 (0 - 2) | - | - |
| Serum RF IgM, median (IQR) | 0 (0 - 3.24) | - | - |
| Serum RF IgA, median (IQR) | 0 (0 - 0) | - | - |
| Average duration in the study, Days (Min, Max) | - | 354.7 (89 - 557) | - |

<sup>1</sup>*P* values were calculated using the paired Wilcoxon rank-sum test for continuous variables. *P* values were calculated using the Fisher exact test for discrete variables.

## Notes

### Summary of Updates

The abstract has been updated to include a link to data visualization tools.

